# Sensory perception of fluctuating light in Arabidopsis

**DOI:** 10.1101/2024.02.21.581400

**Authors:** Antonela Belmonte, Nicolas Tissot, Andrés Rabinovich, Edmundo L. Ploschuk, Carlos D. Crocco, Roman Ulm, Jorge J. Casal

## Abstract

When exposed to shade from neighbours, competitive plants modify their growth patterns to improve access to light. In dense plant stands, ranging from forests to humid grasslands and crops, shade is interrupted by sunflecks penetrating the canopy. Relatively infrequent, minute-scale interruptions can significantly contribute to the daily light input. However, given the short duration and the time gap between these low frequency sunflecks (LFS), whether plants can sense them was unknown. Here we show that phytochrome B (phyB), cryptochrome 1 (cry1), cry2, and UV RESISTANCE LOCUS 8 (UVR8) cooperatively perceive LFS to reduce hypocotyl growth in *Arabidopsis thaliana*. LFS also enhanced the expression of photosynthetic and photo-protective genes and initiated pre-emptive acclimation to water restriction. Repeated LFS increased the nuclear abundance of cry1 and UVR8. This positive feedback enhanced the sensitivity to subsequent LFS and even to the shade between LFS. LFS reduced the nuclear abundance of the growth regulator PHYTOCHROME INTERACTING FACTOR 4 (PIF4), which only slowly recovered upon return to shade, further amplifying the signal. The dynamics of the photo-sensory system under fluctuating light helps adjust plants to the prevailing conditions.

## Introduction

Plants require light for photosynthesis but due to seasonal and daily rhythms in income and the interference caused by neighbouring vegetation, the levels of this resource are seldom optimal. Excessive light can damage the photosynthetic apparatus and limiting light impairs the synthesis of carbohydrates required to sustain plant life (Tikkanen *et al*., 2012; Demarsy *et al*., 2018). To mitigate the impact of shade, plants use a blend of acclimation and shade-avoidance responses, in proportions that depend on the strategy of the species. During acclimation to shade, the photosynthetic apparatus gains efficiency to work at low irradiances (Vialet-Chabrand *et al*., 2017). During shade avoidance, plants differentially affect the growth of their organs to place the leaves higher within the canopy and reach less attenuated light (Casal & Fankhauser, 2023).

Under sunlight, the phytochrome B (phyB) (Burgie *et al*., 2021) and cryptochrome 1 (cry1) (Wang & Lin, 2020) sensory receptors are active and repress PHYTOCHROME INTERACTING FACTORs (PIFs) (Pham *et al*., 2018). Under shade, the activities of phyB and cry1 decrease and PIF transcription factors trigger shade avoidance responses (Lorrain *et al*., 2008; Li *et al*., 2012). For instance, in young seedlings of Arabidopsis exposed to neighbour cues, PIF4, PIF5 and PIF7 promote hypocotyl growth by increasing the synthesis of the growth hormone auxin in the cotyledons and auxin sensitivity in the hypocotyl itself (Li *et al*., 2012; Hornitschek *et al*., 2012). Furthermore, phyB and cry1 repress the activity of the E3 ubiquitin ligase CONSTITUTIVELY PHOTOMORPHOGENIC1 (COP1) (Podolec & Ulm, 2018; Ponnu & Hoecker, 2021). Under shade, COP1 increases its nuclear abundance and targets to degradation negative transcriptional regulators of PIFs (Pacín *et al*., 2013, 2016; Blanco-Touriñán *et al*., 2020). COP1 also protects BRI1-EMS-SUPPRESSOR 1 (BES1) in the hypocotyl, a transcription factor that contributes locally to the promotion of hypocotyl growth (Costigliolo Rojas *et al*., 2022).

The degree of shade received by plant tissues is dynamic but our understanding of how the photo-sensory system of plants interprets light /shade signals with different frequencies or temporal stimulation patterns is only incipient (Smith & Berry, 2013; Sellaro *et al*., 2024). The growth of a given plant relative to that of its neighbour competitors affects the degree of shade to which this plant is exposed in the range of days. Solar elevation affects shade in a range that goes from hours to minutes because direct light can penetrate the canopies through large gaps defined by distant plants and micro-gaps created by the fine weave of leaves at nearly random positions, and the movement of the foliage by wind causes fluctuations of shade in the order of seconds (Kaiser *et al*., 2017; Durand *et al*., 2021). Sun patches (episodes of >8 min duration) and sun gaps (episodes of >2 h duration) reduce shade avoidance, in a process that involves the activation of phyB and cry1, but also of the UV-B photoreceptor UV RESISTANCE LOCUS 8 (UVR8) (Podolec *et al*., 2021a), phyA and cry2 (Sellaro *et al*., 2011; Moriconi *et al*., 2018). Sunflecks have been defined as sun episodes of less than 8 min duration and irradiance levels not reaching those observed above the canopy (Smith & Berry, 2013). Very frequent sunflecks with a short duration of a few seconds predictably establish intermediate levels of phyB and cry1 activity causing intermediate degrees of shade avoidance. However, whether low-frequency shade interruptions for 1-2 minutes affect shade avoidance is still a conundrum. In fact, the promotion of hypocotyl growth initiated by neighbour cues can take approximately 1 h before showing partial reversal after seedling exposure to light (Cole *et al*., 2011). This is an important question, because sunflecks longer than 1 min, although relative infrequent compared to those of a few seconds of duration, are those that have the highest contribution to photosynthesis (Durand *et al*., 2021). This uncertainty undermines one of the fundamental concepts in our understanding of shade-avoidance responses, i.e., that their magnitude inversely relates to the availability of light for photosynthesis, incipient when the plants perceive the risk of future shade and maximal under deep shade, where light is severely limiting. Here we show that Arabidopsis seedlings do respond to low frequency sunflecks (LFS, 2 min duration, separated by 6 min shade) via mechanisms that depend on the dynamics of the signalling components.

## Materials and methods

### Plant material

We used seedlings of *Arabidopsis thaliana* of the Columbia (Col) accession, and the mutant and transgenic lines listed in Table S1. Seeds were sown on 1% agar-water in clear plastic boxes (40 mm x 33 mm x 15 mm) lidded with a transparent film (Rolopac, 0.025-mm thick) and stratified 2–3 d at 5°C in darkness.

### Generation of the phyA phyB cry1 cry2 amiR-uvr8 mutant

The *phyA phyB cry1 cry2 amiR-uvr8* lines were generated with the amiRNA targeting *UVR8* (AT5G63860) transcript as described (Vandenbussche *et al*., 2014). The Agrobacterium strain GV3101 was used to transform *phyA phyB cry1 cry2* plants by the floral-dip method (Clough & Bent, 1998). Transgenic seedlings were selected on substrate containing glufosinate ammonium. UVR8 protein abundance was determined by gel protein blot analysis in the T3 generation and transgenic phenotypes were confirmed in the T4 generation (Fig. S1).

### Light conditions

The boxes with seedlings were placed with the agar oriented vertically under the simulated shade condition (control) at 20°C. A photoperiod of 10 h, a red/far red ratio of 0.2 and a photosynthetically active radiation of 27 μmol m^-2^ s^-1^ was provided by a mixture of white light LED lamps (7 W, OSRAM) and halogen lamps (70 W, OSRAM) in combination with a green filter (no. 089; LEE Filters, Hampshire, UK). In the basic protocol, one hour after the beginning of the 4^th^ photoperiod, some of the boxes containing seedlings were transferred to the white light (WL) or white light plus UV-B (WL+UV-B) LFS (2 min, followed by 6 min shade) during 3 h (Fig. S2a-b). WL LFS of 124 μmol m^-2^ s^-1^ (400-700 nm) were provided by white light LED lamps (7 W, OSRAM) placed under the green filter. UV-B with a peak at 311 nm, 2.3 μmol m^-2^ s^-1^ was provided by a narrowband lamp (PLL/PL-S, Phillips). The spectra of the light sources and the resulting treatments are shown in Figure S2c-d. In the experiments to analyse the sensitivity to blue light, we used a red filter (no. 130; LEE Filters, Hampshire, UK) to cut off the 6.5 μmol m^-2^ s^-1^ of blue light present under the simulated shade condition.

### The choice of LFS duration, frequency and spectral composition

The primary aim of this work is to investigate the sensory perception of sunflecks that make significant contribution to photosynthesis but are infrequent and therefore, less likely to potentiate with subsequent light exposures. We based the choice of duration and frequency on data from soybean crops (Pearcy *et al*., 1990). We selected a duration of 2 min, because these relatively long sunflecks require larger gaps in the canopy, which allow the penetration of stronger photosynthetic light and are among those that contribute more strongly to photosynthesis (Pearcy *et al*., 1990). We selected 6 min of shade between successive sunflecks because we wanted to minimise the probability of carryover from one sunfleck to the next, and taking into consideration that only 10% of the sunflecks are normally preceded by shade periods longer than 6.4 min (Pearcy *et al*., 1990). In natural settings, the spectral photon distribution of shade and during sunflecks depends on the specific conditions (Durand & Robson, 2023), but considering general features, simulated shade contained limited amounts of UV-B and photosynthetically-active radiation, which increased during the sunflecks. We deliberately used weaker UV-B than in more conventional UV-B studies to stay within more natural proportions.

### Hypocotyl growth

Twelve seedlings of each genotype were grown per box (one replicate) parallel to the vertical agar surface. The seedlings were photographed (PowerShot; Canon, Tokyo, Japan) 1 h and 4 h after the beginning of the 4^th^ photoperiod (i.e., during the 3 h exposure to LFS) for the basic protocol, 4 h and 6 h after the beginning of the 4^th^ photoperiod (i.e., immediately after the 3 h exposure to LFS) for the analysis of sensitivity to blue light, or 1 h and 4 h after the beginning of the 5^th^ photoperiod (i.e., 1 d after the exposure to the LFS) in the drought experiments (Fig. S2a). Hypocotyl length increments were measured using image processing software (Global Mapper 11) and divided by the duration of the period to obtain the growth rates.

### Confocal microscopy

We obtained confocal fluorescence images with an LSM5 Pascal, LSM710 and LSM510 (Zeiss) laser scanning microscope with a Plan-Apochromat 40x/1.2, Plan-Neofluar 40x/0.75 and 40x/1.30 Oil EC Plan Neofluar DIC respectively lens. For chloroplast visualisation, probes were excited with a He-Ne laser (λ = 543 nm) and fluorescence was detected using an LP560 filter. For visualisation of GFP/YFP fusion proteins, probes were excited with an Argon laser (λ = 488 nm) and fluorescence was detected using BP 505-530. For visualisation of mCherry fusion proteins we used He-Ne laser (λ = 543 nm), LP560 filter and used the spectral detector to separate mCherry signal from chlorophyll autofluorescence. We measured the fluorescence of all the nuclei in each image and divided the sum of these fluorescence values by the number of cells in the image. Therefore, the nuclei with no detectable fluorescence did not contribute to the numerator of the equation but their cells were included in the denominator. To calculate the number of cells we measured the area of five cells and divided the area of the image by the average area of a cell. Images of the selected nuclei were obtained by using the same configuration and 8X magnification. All the measurements were made in ImageJ software (https://imagej.nih.gov/).

### Net carbon dioxide exchange

We used a portable gas-exchange system (Li-Cor 6400; Li-Cor, Lincoln, NE, USA) to measure net CO_2_ exchange (μmol CO_2_ m^−2^ s^−1^). This equipment automatically controls air flow (300 μmol s^−1^) and CO_2_ concentration in the reference cell (CO_2_R, 400 ppm). We cultivated a lawn of seedlings on a nylon mesh placed on top of the agar. Then, we transferred the agar and the membrane containing seedlings to a custom-built cylindrical chamber (20 cm^3^) connected to the gas-exchange system and placed under external light sources that provided 27 μmol m^−2^ s^−1^ of shade light or 2000 μmol m^−2^ s^−1^ of WL.

### Immunoblot analysis

For analysis of UVR8, CHS and actin, total proteins of complete seedlings were extracted in 50 mM Tris, pH 7.6, 150 mM NaCl, 2 mM ethylenediaminetetraacetic acid (EDTA), 10% (v/v) glycerol, 5 mM MgCl_2_, 1% (v/v) Igepal (Sigma), 1% (v/v) protease inhibitor mixture for plant extracts (Sigma), and 10 μM MG132. The concentration was determined using the Bio-Rad Protein Assay Dye Reagent Concentrate according to the manufacturer’s instructions. To determine total UVR8, CHS and actin, samples were boiled. To identify the UVR8 homodimers the boiling step was omitted (Rizzini *et al*., 2011). Proteins were separated by 12% SDS-PAGE and gels were UV-B-irradiated before electrophoretic transfer to polyvinylidene difluoride (PVDF) membrane. Anti-UVR8^426-440^ (1:4000), anti-CHS (Santa Cruz Biotechnology, 1:10000), and anti-Actin (A0480, Sigma-Aldrich, 1:20000) were used as the primary antibodies (Favory *et al*., 2009). Conjugated anti-rabbit and anti-mouse immunoglobulins were used as the secondary antibodies (Agilent Dako, 1:20000). Immunodetection was performed using an ECL Plus Western Detection Kit and revealed with an ImageQuant LAS 4000 mini-CCD camera system (GE Healthcare). ImageJ was used for quantifications of the bands.

### Quantitative real-time PCR

Isolation of total RNA (including DNase treatment) and synthesis of cDNA were conducted with Plant RNeasy kit (Qiagen) and the TaqMan Reverse Transcription Reagents kit (Thermo Fisher Scientific), respectively, following manufacturer’s standard protocols. Each quantitative real-time PCR reaction contained cDNA synthesised with a 1:1 mixture of oligo(dT) primers and random hexamers from 300 ng of total RNA and was performed using a PowerUp SYBR Green Master Mix (Thermo Fisher Scientific). The primers for *CHS* and *PROTEIN PHOSPHATASE 2A SUBUNIT A3 (PP2AA3)* were as reported (Czechowski *et al*., 2005; Xu *et al*., 2016)

### RNAseq experiment

Seedlings were harvested in liquid nitrogen and total RNA was extracted using a Spectrum Plant Total RNA kit (Sigma-Aldrich). The sequencing was made on an Illumina HiSeq 2500 system using 100-bp single-end reads protocol, performed at the iGE3 genomics platform of the University of Geneva. Single-end 100bp reads were processed with TrimGalore version 0.6.6 (TRIMGALORE) to remove residual adaptor sequences at both 5′ and 3′ from raw reads and to filter fragments with a phred score below 20 (default value). FastQC v0.11.5 (FASTQC) was used to examine sequencing files before and after adaptor trimming and quality filtering. Reads were aligned to the Arabidopsis TAIR10 release 52 genome (TAIR) using STAR version 2.7.9a (STAR). ASpli version 2.10.0 (ASPLI) was used for transcript quantification. Gene references were obtained using org.At.tair.db v3.14.0. Data have been deposited in NCBI BioProyect portal as BioProject ID PRJNA1040773.

### PEG treatments

PEG was used to generate water restriction (Verslues *et al*., 2006; Osmolovskaya *et al*., 2018). We added 1.5 ml solutions containing 0 g/l (water control), 400 g/l or 700 g/l of PEG of a molecular weight of 8000 (BIOFROXX, 25322-68-3) per 1 ml of agar 1% substrate, incubated for 24 h and discarded the excess of solution. One hour after the beginning of the 5^th^ photoperiod (i.e., 1 d after the exposure to LFS), we transferred the nylon mesh with seedlings from their original agar to agar equilibrated with the above solutions (Fig. S2a).

### Statistical analyses

We used the boxes of seedlings as biological replicate (i.e., for hypocotyl growth we used the average of 12 seedlings in each box). Hypocotyl growth rate can be modelled as the maximum growth rate divided by one plus the terms corresponding to the inhibitory effects caused by the different photoreceptors and light conditions (Romero-Montepaone *et al*., 2020). Thus, to analyse the contribution of different photoreceptors to the control of hypocotyl growth (Fig. 1d), we used stepwise linear regression with hypocotyl growth rate^-1^ as response variable and the explanatory variables listed in Table S2, from which the stepwise analysis filtered out those that were not significant. For other hypocotyl growth, confocal microscopy, and protein gel blot data we conducted Student’s *t* tests or ANOVA followed by Tukey multiple comparison tests (GraphPad Prism 7 software, San Diego, California). For the analysis of the transcriptome, Voom normalisation of RNAseq data (Law *et al*., 2014) was conducted on EdgeR v3.42.4(R Core Team, 2017). Normalised data were subjected to two-way ANOVA (LFS conditions and genotypes as main effects) in RStudio v4.1.3. We used a *q*-value (Storey & Tibshirani, 2003) <0.01 cut-off to identify a group of genes that responded to LFS (main effect or interaction with genotype). These genes were classified according to their response to genotype using a p-value <0.05. This second threshold was more permissive to minimise the inclusion of genes apparently affected by the photoreceptors within the clusters described as unaffected by the photoreceptors. The heatmaps were generated with RColorBrewer v1.1-2, pheatmap v1.0.12. Overrepresented Gene Ontology (GO) terms were identified by using PANTHER v18.0 (Thomas *et al*., 2003). The graphs of overrepresented GO terms linked by functional roles were generated by using the visNetwork package (Almende *et al*., 2019), including broad categories with at least 500 genes and their associated categories sharing at least 20% of common features.

**Fig. 1.**
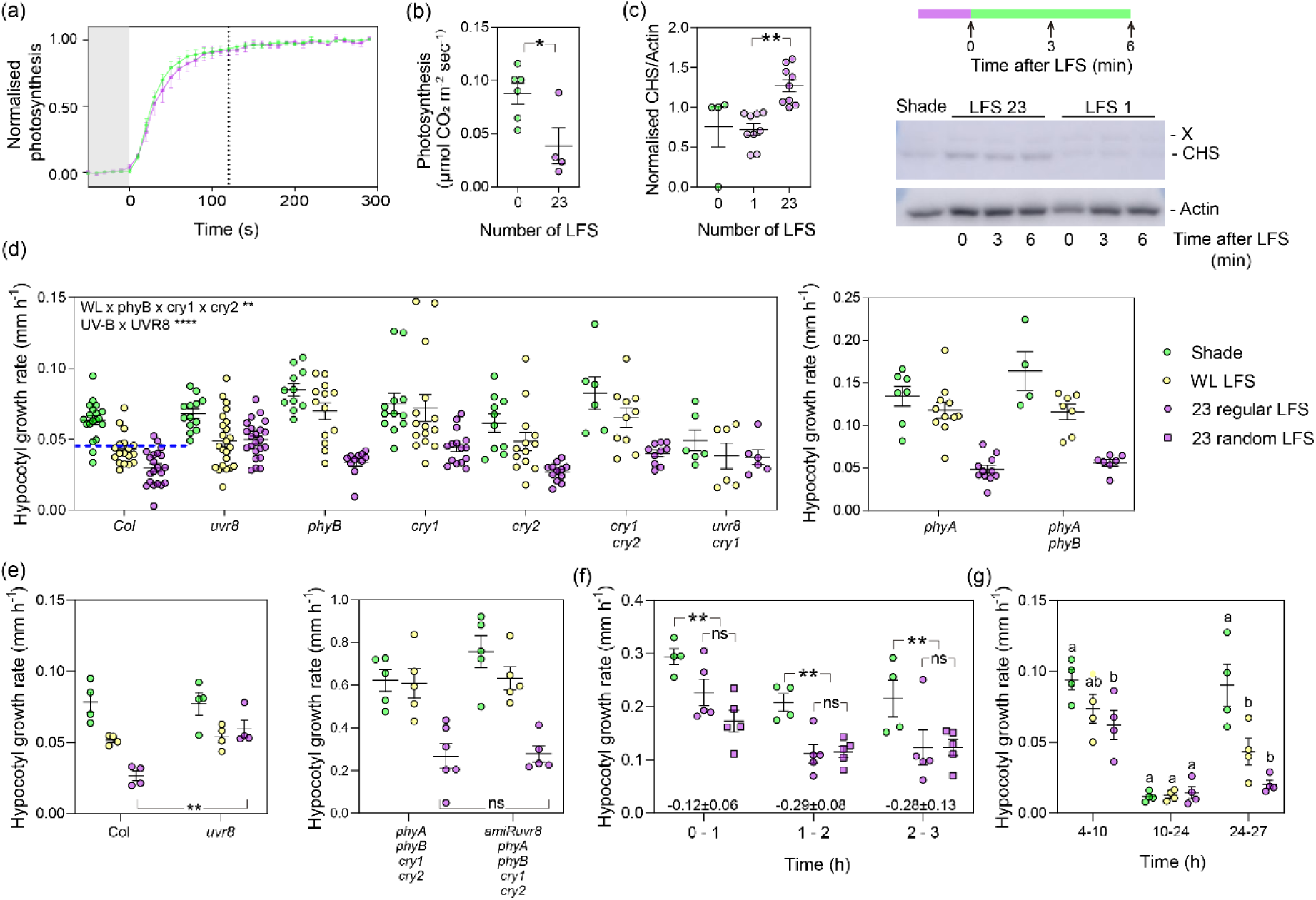
Arabidopsis seedlings grown under shade respond to LFS. (a) Photosynthesis of Arabidopsis seedlings approaches maximum rates within two minutes of exposure to white light (2000 µmol m^-2^ s^-1^). Photosynthesis normalised by setting shade rates to 0 and maximum rates to 1. The seedlings were grown under shade or WL+UV-B LFS 3 h day^-1^ during days 4 to 6 and photosynthesis was measured on day 7 (Fig. S2a). (b) Actual maximum rates of photosynthesis of the seedlings depicted in (a). (c) LFS increase the abundance of CHS protein compared to shade, but a single LFS is ineffective. Left panel, quantification of CHS protein levels relative to actin; right panel, representative anti-CHS and anti-actin protein gel blot analysis (“x” unspecific band). The seedlings were exposed to 1 LFS followed by 172 min shade or 23 WL+UV-B LFS (3 h) before harvest. (d-e) Rate of hypocotyl growth in response to LFS. The dashed line (d, Col) assumes no growth during the 2 min of each LFS, shade growth rates between LFS and equal lags for both transitions. Hypocotyl growth was measured during the 3 h of exposure to WL or WL+UV-B LFS. LFS started 1 h after the beginning of the shade photoperiod (Fig. S2a). (f) Kinetics of hypocotyl growth rate of Col seedlings during the 3 h of exposure to 23 LFS separated either by regular shade intervals (6 min) or random shade intervals (2, 4, 6, 8 or 10 min assigned randomly). Percent inhibition caused by regular LFS is indicated for each period at the base of the figure. (g) Hypocotyl growth rate of Col seedlings after the 3 h of exposure to 23 LFS (treatment: 1-4 h, night: 10-24 h). Data are means ±SE and individual values (b-g) of 4-21 biological replicates (in d-g, each biological replicate involves 12 seedlings). In (b-c) and (e-g), asterisks indicate significant differences (*, P <0.05; **, P <0.01, ns, not significant) in *t* (b-c, e) and Dunnet’s multiple comparisons (f-g) tests. In (d), multiple regression analysis indicates significant interaction between the WL LFS phyB, cry1 and cry2 and between the addition of UV-B and UVR8 and a basal effect of phyA (Table S2).

## Results

### LFS are input and stress for the photosynthetic apparatus

Since the function of shade avoidance is to increase the chances to capture more light for photosynthesis, we investigated the effect of LFS on the photosynthetic apparatus to provide context to the subsequent shade-avoidance experiments. We grew seedlings of Arabidopsis under simulated shade or under simulated shade interrupted by LFS providing white light plus UV-B radiation (WL+UV-B), 3 h day^-1^ for 3 days (see protocol and spectral distribution in Fig. S2). Figure 1a shows the kinetics of the rate of photosynthesis in seedlings exposed to 2000 µmol m^-2^ s^-1^ of photosynthetically active radiation (to focus on the kinetics, data are normalised using 0 for the values under shade and 1 for maximum photosynthesis). By the end of the first 2 min, the rate of photosynthesis had already reached values close to the maximum. However, the maximum rate of photosynthesis was higher in shade-grown seedlings than in the seedlings pre-treated with UV-B LFS (Fig. 1b). Therefore, sunflecks of 2 min offer enough time to leverage the light input but reduce the photosynthetic efficiency.

### Arabidopsis seedlings perceive LFS

A single LFS was not effective, but 23 LFS increased the abundance of CHALCONE SYNTHASE (CHS) protein, an enzyme involved in the synthesis of photoprotective pigments (Fig. 1c). The difference between 1 and 23 LFS was associated with differences in the expression of the *CHS* gene (*CHS* expression relative to *PP2AA3*, mean ±SE, n=4, 1 LFS: 0.4 ±0.1; 23 LFS: 2.6 ±0.5, P= 0.005). In Arabidopsis, shade stimulates the growth of the hypocotyl to place the cotyledons at higher, better lit positions within the canopy. Another way to mitigate the potential damage of the photosynthetic apparatus would be by reducing hypocotyl growth and hence foliage exposure to higher light inputs. Seedlings grown under shade were either exposed to LFS or remained as shade controls for 3 h, whilst hypocotyl growth was measured (Fig. S2). LFS inhibited the growth of the hypocotyl (Fig. 1d). LFS containing white light and no UV-B (WL LFS) were per se effective, but WL+UV-B LFS caused a stronger inhibition.

### Synergic co-action among photoreceptors in the response to LFS

We used different photoreceptor mutants to investigate their contribution to the control of hypocotyl growth under shade and in response to the WL and UV-B components of the LFS. The statistical analysis selected the interaction among WL, phyB, cry1 and cry2, the interaction between UV-B and UVR8, and a basal effect of phyA under shade as explanatory variables (Fig. 1d, Table S2). The analysis filtered out the simple interactions between WL and anyone of these photoreceptors alone, indicating that the response to WL LFS required phyB, cry1 and cry2, and was not significant if one of them was lacking. This is a case where the contribution of a photoreceptor to a physiological response is enhanced by the activation of other photoreceptor(s); a phenomenon called synergic co-action or responsivity amplification (Mohr, 1994; Casal, 2000).

To investigate whether the photoreceptors that are active under WL LFS (phyB, cry1, cry2) or shade (phyA) can contribute to the UVR8-mediated response, we generated *phyA phyB cry1 cry2 amiR-uvr8* seedlings, bearing a severe reduction of UVR8 (Fig. S1). In the wild-type background, UVR8 inhibited hypocotyl growth in seedlings exposed to WL+ UV-B LFS (Col and *uvr8* are different under this condition); conversely, in the *phyA phyB cry1 cry2* mutant background UVR8 was ineffective to inhibit hypocotyl growth (note no difference between the quadruple mutant and the *phyA phyB cry1 cry2 amiR-uvr8* line, Fig. 1e). This lack of detectable UVR8-mediated effects occurred despite the inhibition caused by UV-B, likely through the induction of damage in the sensitive *phyA phyB cry1 cry2* background. Taken together, these observations indicate that under LFS there is synergic co-action between UVR8 and other photoreceptors (Mohr, 1994; Boccalandro *et al*., 2001). Synergic co-action is typically conditional; it occurs when the light input is suboptimal (as it is the case under LFS) but not under stronger light inputs, when the photoreceptors tend to be redundant (Casal & Mazzella, 1998; Sellaro *et al*., 2024). In accordance with this concept, under continuous WL+UV-B, *phyA phyB cry1 cry2* seedlings were significantly shorter than *phyA phyB cry1 cry2 amiR-uvr8* seedlings, demonstrating a clear action of UVR8 in the absence of the other photoreceptors when the light input is stronger (Fig. S1). Under continuous light, the *uvr8* phenotype was stronger in the *phyA phyB cry1 cry2* than in the Col background (Fig. S1), evidencing some degree of redundancy.

### LFS have persistent effects on hypocotyl growth

Although to facilitate the analysis we used LFS separated by regular shade intervals, in nature LFS occur at random, irregular intervals. Thus, we compared regular LFS with 2 min LFS randomly distributed during the treatment period (shade intervals between successive LFS were 2, 4, 6, 8 or 10 min). The kinetics of growth inhibition was not significantly different (Fig. 1f), validating the use of the regular LFS.

The effect of each LFS cannot be resolved in real time but it is possible to calculate the growth rate assuming that after an initial lag, the WL+UV-B LFS completely ceases growth for 2 min, and after the end of the LFS and a second lag similar in duration to the first lag, elongation growth recovers the rates observed in shade controls (dashed line in Figure 1d). The actual inhibitory effect of LFS is stronger than calculated. Since the strength of inhibition used for the calculation is already maximal during the LFS (no growth), this result indicates that the lag for growth recovery must be longer than the lag to establish growth inhibition, i.e., LFS have a persistent effect. Of the 23 LFS displayed at regular shade intervals, eight were given during the first hour and already caused a significant inhibition of hypocotyl growth. However, as further LFS are experienced during the second and third hours the inhibitory effects become stronger (Fig. 1f), indicating that early LFS have effects beyond their own duration. Hypocotyl growth did not fully recover under shade after the termination of the LFS treatment (4-10 h, Fig. 1g). Of note, the persistent effect was not evident during the subsequent night but reappeared the following morning under shade (10-24 and 24-27 h, Fig. 1g), suggesting that it might involve enhanced photoreceptor activity and sensitivity to light.

### LFS do not affect the nuclear bodies of phyB

WL LFS did not affect the nuclear abundance of phyB (Fig. 2a), which is consistent with the relatively slow migration of active phyB to the nucleus and the fact that phyB does not leave the nucleus upon phototransformation to the inactive form (Klose *et al*., 2015). The brightness of the phyB nuclear bodies correlates with phyB activity (Van Buskirk *et al*., 2014; Chen *et al*., 2022) but despite the physiological role of phyB (Fig. 1d), WL LFS did not affect the brightness, size, or number of phyB nuclear bodies (Fig. 2b and Fig. S3).

**Fig. 2.**
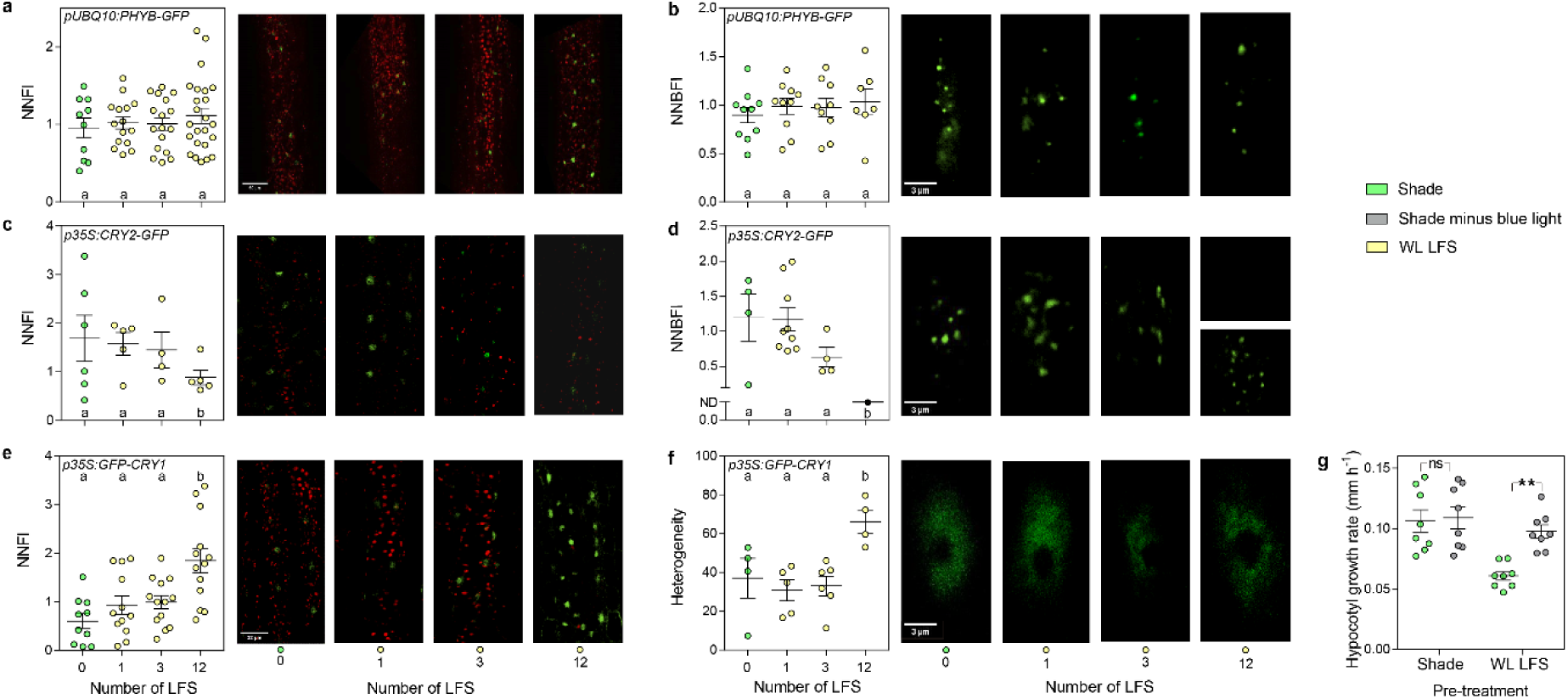
LFS enhance cry1 nuclear abundance and the sensitivity to blue light. (a-b) The normalised nuclear fluorescence intensity (NNFI, a) and the normalised nuclear body fluorescence intensity (NNBFI, b) of phyB-GFP do not respond to the LFS. (c-d), The NNFI (c) and NNBFI (d) of cry2 decrease in response to LFS. After 12 LFS the nuclear bodies were not detected with the standard gain used for the confocal microscope, but they were detectable by increasing the gain (d, upper and lower photographs, respectively). (e-f) The NNFI (e) and the heterogeneity of the NNFI (f) of cry1 increase in response to LFS. The heterogeneity was calculated as the ratio between the highest and lowest fluorescence regions of a fixed area within the nucleus. (g) LFS increase the hypocotyl growth inhibition caused by weak blue light present under subsequent shade conditions. Confocal images were taken at the end of the indicated LFS or under shade (simultaneously with the plants exposed to LFS). LFS started 1 h after the beginning of the shade photoperiod (Fig. S2a). Data are means ±SE and individual values of 4-24 biological replicates and representative confocal images are shown. Significant differences in Tukey’s multiple tests indicated by different letters (P < 0.05) or asterisks (**, P <0.01, ns, not significant).

### LFS decrease cry2 nuclear abundance

cry2 forms nuclear bodies by phase separation (Wang *et al*., 2021) and, consistently with its light lability (Lin *et al*., 1998), repeated WL LFS reduced nuclear fluorescence and nuclear body fluorescence driven by cry2-GFP, which was still detectable after 12 LFS (Fig. 2c-d, S4a). This residual cry2 could explain the contribution to the hypocotyl growth response to LFS.

### LFS increase cry1 abundance and sensitivity to blue light

WL LFS significantly increased the nuclear abundance of cry1-GFP (Fig. 2e), which was paralleled by an increase in total cellular fluorescence (Fig. S4b). cry1 did not form nuclear condensates upon exposure to WL LFS; however, repeated LFS generated the appearance of areas with more cry1 and areas with less cry1 within the nucleus (Fig. 2f), which is suggestive of a partial process of condensation. In fact, 1 h exposure to white light does lead to the generation of visible nuclear bodies (Fig. S4c, (Liu *et al*., 2022)).

Increasing the number of WL LFS did not enhance the effect of each LFS, as might be expected from the increased cry1 nuclear levels (Fig. 2e). Therefore, in additional experiments, we measured hypocotyl growth under shade conditions after (rather than during) the 3 h of exposure to WL LFS to test the possibility that LFS enhance the sensitivity to shade (instead of the sensitivity to WL LFS). We included control seedlings not exposed to WL LFS (shade control) and for both groups we measured growth in seedlings exposed to shade devoid of its blue-light component. The seedlings always grown under shade did not elongate differently under further shade with or without its blue-light component (Fig 2g). However, the seedlings that had been pre-treated with WL LFS did respond to blue light present under shade. Therefore, plants exposed to WL LFS acquire enhanced sensitivity to blue light, consistently with their increased nuclear levels of cry1.

### LFS increase the nuclear abundance of UVR8 and the response to UV-B

Repeated WL+UV-B LFS elevated the nuclear abundance of YFP-UVR8 (Fig. 3a). We used protein gel blot analysis of heat-denatured samples to quantify total UVR8 protein and, consistently with previous reports showing no effects of prolonged UV-B, we observed no response to LFS (Fig. 3b) (Kaiserli & Jenkins, 2007; Favory *et al*., 2009; Yin *et al*., 2016). UV-B causes the monomerization of the UVR8 homodimer (Rizzini *et al*., 2011). In protein blots using non heat-denatured conditions, we observed a dimer / monomer ratio consistent with the presence of UV-B under our shade conditions (Findlay & Jenkins, 2016; Moriconi *et al*., 2018; Liao *et al*., 2020). One or 23 LFS similarly reduced the dimer / monomer ratio (0 min after LFS in Fig. 3c, S5). Depending on the conditions, after prolonged exposure to UV-B, 50 % re-dimerization in vivo can take from 18 to 60 min (Heilmann & Jenkins, 2013; Heijde & Ulm, 2013), (Fig. S6). Consistently with these rates, we did not detect significant UVR8 re-dimerization during the shade period after 23 LFS (Fig. 3c). Surprisingly, after the first LFS substantial re-dimerization occurred during the subsequent 6 min shade, suggesting that a brief exposure to UV-B is not sufficient to stabilise the monomer in vivo.

**Fig. 3.**
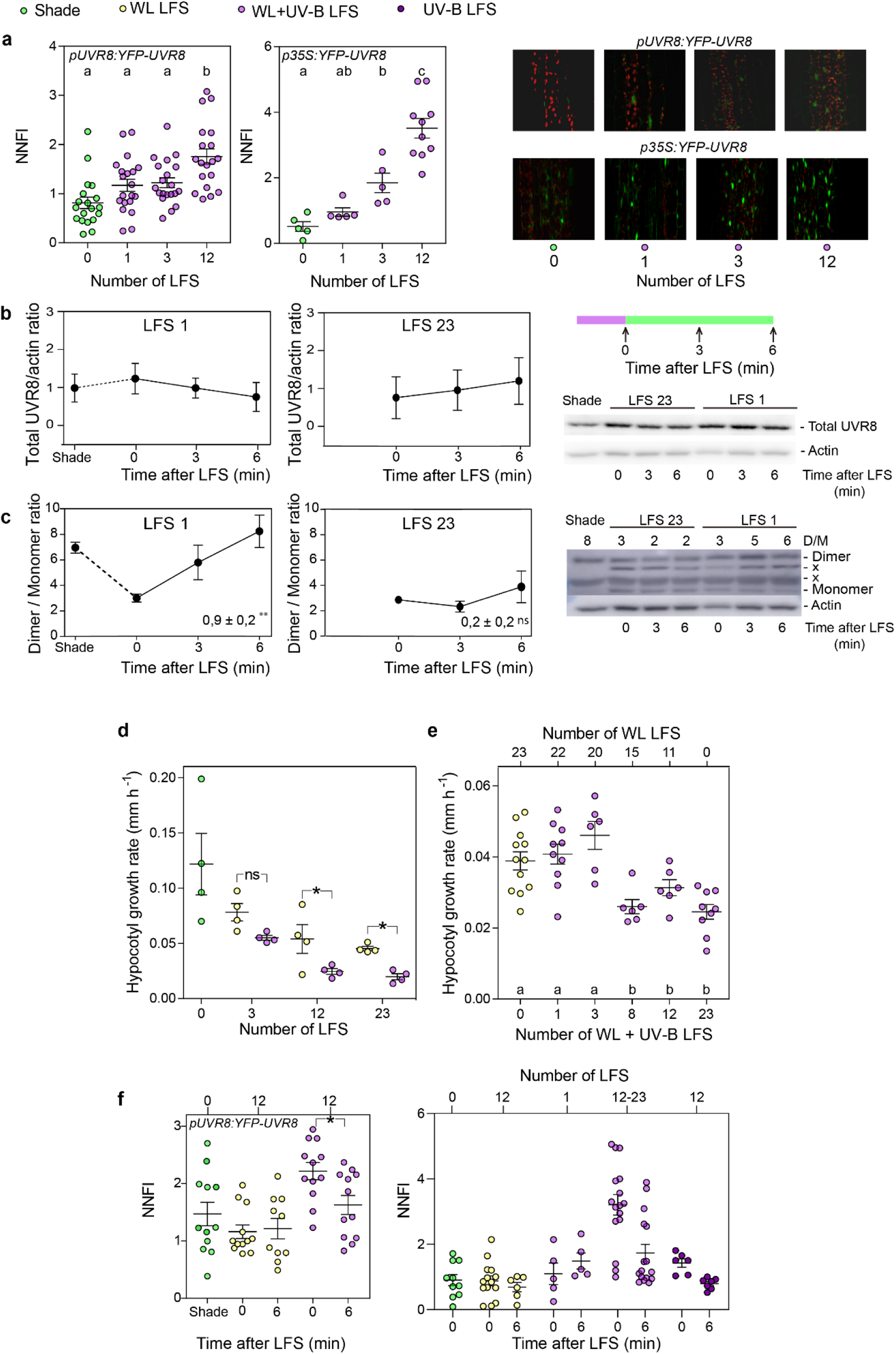
LFS enhance UVR8 nuclear abundance and the sensitivity to UV-B. (a) WL+UV-B LFS increase the normalised nuclear fluorescence intensity (NNFI) of YFP-UVR8. Confocal images were taken immediately after the end of the indicated LFS and in shade controls. (b) LFS do not affect UVR8 protein abundance. Seedlings were harvested for protein extraction at the indicated times after a LFS simultaneously with shade controls. (c) LFS slowed down UVR8 re-dimerization. The slopes ±SE of the recovery lines and their significance are indicated. (d-e) UV-B becomes effective only after repeated WL+UV-B LFS. Hypocotyl growth was measured during the 3 h in shade controls and in seedlings exposed to increasing number of WL or WL+UV-B LFS (d), or a fixed number of LFS (23) but either of WL or WL+UV-B (e, UV-B was evenly distributed among the 23 LFS by changing the frequency to modify the number). (f) UVR8 partially leaves the nucleus during the shade periods after WL+UV-B LFS and UV-B LFS without WL are less effective to induce UVR8 nuclear accumulation. Confocal images were taken at the indicated times after a LFS or under shade (simultaneously with the plants exposed to LFS). LFS started 1 h after the beginning of the shade photoperiod (Fig. S2a-b). Data are means ±SE and individual values of 4-20 biological replicates and representative confocal or protein gel blot images are shown (“x” unspecific bands, D/M: dimer / monomer ratio). Significant differences in Tukey’s multiple tests indicated by different letters (P < 0.05) or asterisks (*, P <0.05; **, P <0.01, ns, not significant).

To investigate the consequences of increased UVR8 nuclear levels, we analysed the impact of increasing the number of LFS on hypocotyl growth. Three LFS of WL or WL+UV-B were not differentially effective but 12 or 23 LFS of WL+UV-B were significantly more effective than a similar number of WL LFS (Fig. 3d). We also exposed the seedlings to 23 WL LFS and modified the number of them that also contained UV-B. Up to 3 LFS containing UV-B caused no difference when compared to the WL LFS alone, but 8 or 23 LFS with UV-B revealed the specific effect of adding UV-B (Fig 3e). These experiments indicate that UV-B present in early LFS sensitise the seedlings to UV-B present in later LFS, likely because they increase UVR8 nuclear levels (Fig. 3a). Whilst this dynamic change in sensitivity enhances the response to WL+UV-B LFS, the constitutively high sensitivity of the *uvr8-17D* and *rup1 rup2* mutants (impaired in UVR8 dimerization (Gruber *et al*., 2010; Heijde & Ulm, 2013; Podolec *et al*., 2021b)) impeded the response to LFS, because the weak UV-B present under shade (Fig. S2) saturated hypocotyl growth inhibition (Fig. S7).

Following prolonged UV-B exposures, UVR8 leaves the nucleus after several hours of recovery without UV-B (Fang *et al*., 2022). Nuclear levels of UVR8 were elevated at the end of the 12^th^ or 23^rd^ 2 min LFS (Figs 3a, f), and decreased during the subsequent 6 min back under shade, indicating a rapid translocation to the cytosol (Fig. 3f). This rapid translocation occurred as monomer because re-dimerization was not detectable during the 6 min under shade (see 23 LFS in Fig. 3c). Exposure to UV-B LFS without WL induced weaker nuclear accumulation of UVR8 than WL+UV-B (Fig. 3f), suggesting that WL photoreceptors facilitate UV-B-induced UVR8 nuclear accumulation under LFS.

### LFS decrease the nuclear abundance of PIF4

We investigated the contribution of signalling components that act downstream the photo-sensory receptors in the control of hypocotyl growth. The *pif4*, *cop1,* and *bes1* mutants failed to respond significantly to the LFS whilst the *hy5* mutant did respond (Fig. 4a-d). Prolonged exposures to UV-B reduce the levels of PIF4 and PIF5 (Hayes *et al*., 2014; Sharma *et al*., 2019; Tavridou *et al*., 2020b,a) but one LFS was not effective (Fig. 4e). Exposure to three LFS of WL or WL+UV-B similarly decreased the nuclear levels of PIF4-GFP measured by confocal microscopy. Twelve LFS of WL+UV-B caused a stronger reduction of PIF4-GFP levels than a similar number of WL LFS (Fig. 4e). Thus, repeated LFS reduced nuclear levels of PIF4-GFP, resembling their effect on hypocotyl growth (see Fig. 3d-e).

**Fig. 4.**
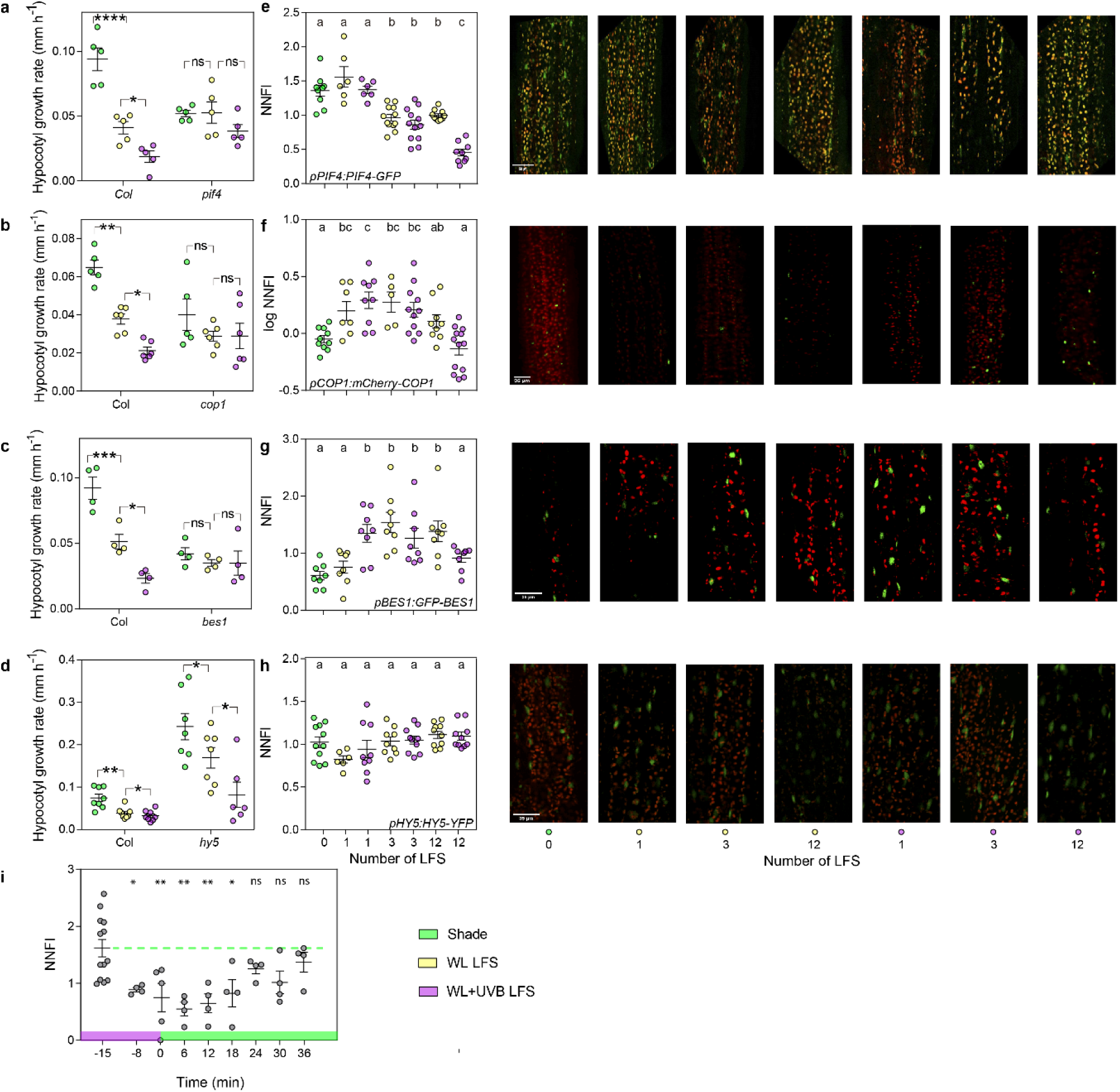
LFS reduce PIF4 nuclear abundance. (a-d) The hypocotyl growth response to the LFS requires PIF4 (a), COP1 (b), and BES1 (c) but not HY5 (d). (e-h) LFS reduce PIF4 (e) normalised nuclear fluorescence intensity (NNFI) and transiently increase the NNFI of COP1 (f) and BES1 (g), without affecting the NNFI of HY5 (h). Hypocotyl growth was measured during the 3 h of exposure to LFS. Confocal images were taken at the end of the indicated LFS and in shade controls. LFS started 1 h after the beginning of the shade photoperiod (Fig. S2a). (i) The decay in PIF4 abundance caused by UV-B is faster than its recovery after returning to shade (shade-grown seedlings received a single 15 min exposure to UV-B followed by shade, the dashed line extends the mean before UV-B to aid the visualisation of the recovery). Data are means ±SE and individual values of 5-13 biological replicates, and representative confocal or protein gel blot images. Significant differences in Tukey’s multiple tests indicated by different letters (P < 0.05) or asterisks (*, P <0.05, **, P <0.01, ***, P <0.001 ****, P <0.0001, ns, not significant). In (i), means are compared to the shade control.

In hypocotyl cells, the nuclear levels of COP1 (Pacín *et al*., 2013, 2014; Moriconi *et al*., 2018) and BES1 (protected from degradation by COP1) (Costigliolo Rojas *et al*., 2022) are lower under sustained white light than under shade; however, the earliest LFS increased the nuclear levels of both mCherry-COP1 and GFP-BES1 (Fig. 4f-g). This could result from COP1 stabilisation due to interaction with photoreceptors (Oravecz *et al*., 2006; Favory *et al*., 2009; Heijde & Ulm, 2013; Huang *et al*., 2013, 2014), together with insufficient time to leave the nucleus during LFS. HY5-YFP did not change its nuclear levels in response to the LFS (Fig. 4h).

Given the ability of PIF4 levels to respond to the LFS, we compared the kinetics of PIF4 decay under WL+UV-B and recovery after termination of the exposure to this condition. We transferred seedlings grown under shade to WL+UV-B and 7 min (from -15 to -8 min) of WL+UV-B caused a significant reduction of nuclear PIF4 levels. However, 18 min after the shift back to shade, PIF4 levels were as low as observed at the end of WL+UV-B exposure (Fig. 4i). Thus, the recovery of nuclear PIF4 is much slower than its decay under WL+UV-B.

### Binding targets of PIF4 are overrepresented among the genes that respond to LFS

To explore the biological significance of the perception of fluctuating light, we analysed the transcriptome of wild type, *cry1 cry2* and *uvr8* mutant seedlings grown under shade and exposed to WL or WL+UV-B LFS. We identified 6400 genes with expression significantly affected by the LFS and genotype. These genes grouped on two major clusters (Fig. 5a), one with expression promoted by the LFS (cluster 1, 3245 genes) and the other with the opposite response (cluster 2, 3155 genes). The responses to WL LFS were reduced in *cry1 cry2* and the additional effects of UV-B were reduced in *uvr8* (Fig. 5a, S8-9). Consistently with the effects of LFS on PIF4 nuclear abundance, the binding targets of PIF4 (Oh *et al*., 2012) were strongly overrepresented among the genes with expression reduced by LFS (cluster 2, 28%, compared to 15% among the genes that did not respond to the LFS, Chi square test with Yates correction: P <0.00001). Although weakly, PIF4 binding target genes were also overrepresented among the genes with expression enhanced by the LFS (cluster 1, 18% compared to 15% of the control group, P <0.00001). Other, smaller gene clusters showed significant effects of LFS (*q* <0.001) and reduced effects of WL LFS in *cry1 cry2* (clusters 3-6), reduced effects of WL+UV-B in *uvr8* (clusters 7-10), or not significant effects of the *cry1 cry2* or *uvr8* mutations (Fig. S10-12).

**Fig. 5.**
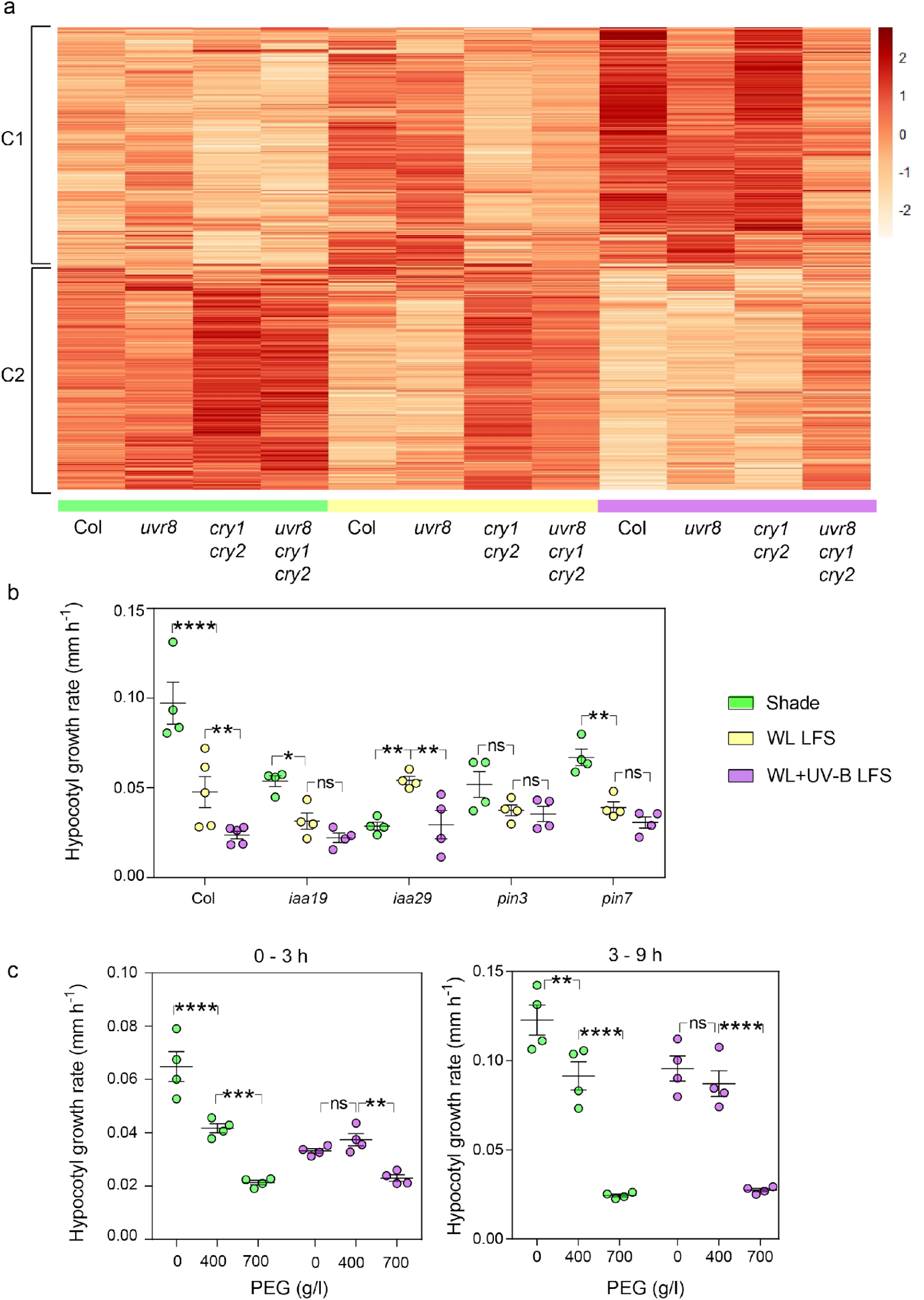
Transcriptome responses to LFS point to the role of auxin in the growth response and uncover acclimation to water deprivation. (a) The heatmap of genes that respond to LFS and genotype defines two major gene clusters, one with expression enhanced by LFS (C1) and the other with expression reduced by LFS (C2). The heatmap is based on three biological replicates harvested on day 4, at the end of the 3 h of exposure to LFS, which started 1 h after the beginning of the shade photoperiod (Fig. S2a). (b) Mutations at the auxin transport genes *PIN3* and *PIN7* and auxin perception genes *IAA19* and *IAA29* impair the hypocotyl growth response to LFS. (c) Plants exposed to LFS are less sensitive to water deprivation. Hypocotyl growth was measured on day 4, during the 3 h of exposure to LFS (b), or on day 5, between 0-3 h and 3-9 h after the transfer to PEG (c, Fig. S2a). In (b-c) data are means ±SE and individual values four biological replicates. Asterisks indicate significant differences in Tukey’s multiple tests (*, P <0.05, **, P <0.01, ****, P <0.0001, ns, not significant).

### LFS reduce the expression of auxin-related genes

In cluster 2, the most significant overrepresented GO terms were “growth” and “small molecule metabolic process” (Fischer test corrected for false discovery rates, P= 1.48E-13 and 6.34E-14, respectively, Fig S8). Both blue light perceived by cry1 and UV-B perceived by UVR8 reduce the expression of auxin responsive genes when added to plants exposed to shade (Hayes *et al*., 2014; De Wit *et al*., 2016; Tavridou *et al*., 2020b). Consistently with those observations, the GO terms “response to auxin”, “auxin transport” and “shade avoidance” were overrepresented in the cluster of genes with expression repressed by LFS (P= 9.98E-04, 1.75E-02 and 4.78E-02 respectively, Fig. S8). These groups include the *IAA19*, *IAA29*, *PIN3* and *PIN7* genes. The *iaa19*, *iaa29*, *pin3* and *pin7* loss-of function mutant showed reduced or distorted hypocotyl growth responses to LFS (Fig. 5b), supporting a role of these changes in gene expression in the physiological response.

### LFS enhance the expression of photosynthesis-related genes

As expected, given the nature of the treatments and the patterns of the *cry1 cry2* and *uvr8* mutants, the GO terms “response to blue light” (P= 3.05E-13) and “response to UV-B” (P= 1.77E-05) were overrepresented among the genes with expression enhanced by the LFS (cluster 1, Fig. S9). The presence of “photosynthesis, dark reaction” (P= 5.45E-05), “regulation of photosynthesis, light reactions” (P= 1.25E-03), “photosynthetic electron transport chain” (P= 3.11E-04), “NADP metabolic process” (P= 4.31E-03), “chloroplast RNA processing” (P= 3.59E-03), “photosystem II assembly” (P= 2.59E-02), “chlorophyll metabolic process” (P= 1.78E-09), “xanthophyll metabolic process” (P= 4.41E-03), “regulation of flavonoid biosynthetic process” (P= 3.06E-06), among other overrepresented GO terms (Fig. S9), is suggestive of a profound acclimation of the photosynthetic apparatus in response to the LFS.

### LFS enhance the expression of water deprivation-related genes

Unexpectedly, “response to water deprivation” was among the overrepresented GO terms in cluster 1 (P= 1.84E-06, Fig. S9). To explore whether these changes could be functionally significant, we investigated the hypocotyl growth response to a reduction in water availability of plants exposed to LFS, compared to their shade controls. For this purpose, we followed the basic protocol, exposing the seedlings to the 3 h of LFS treatments during day 4, but then, on day 5 we transferred seedlings to a substrate containing 400 or 700 g/l of polyethylene glycol (PEG) or to fresh substrate without PEG and measured growth during the first 3 h and the subsequent 6 h, under shade (see protocol in Fig. S2a). In the seedlings that remained under shade during day 4, 400 g/l PEG significantly reduced hypocotyl growth, compared to the water control, and 700 g/l PEG caused a more intense effect (Fig. 5c). Conversely, in the seedlings exposed to LFS during the previous day, 400 g/l PEG were completely ineffective to inhibit growth compared to the water control, although 700 g/l were inhibitory. This indicates that the exposure to LFS renders the seedlings less sensitive to water restriction. It is interesting to note that the water controls showed a persistent effect of the LFS experienced the previous day as all the seedlings of the PEG experiments remained under shade during growth measurements (Fig. 5c).

## Discussion

The threat posed by competitors for light depends on the architecture of the canopy, which affects not only the irradiance and spectral composition of the background diffuse light under shade, but also the frequency of penetration of direct light during sunflecks (Burgess *et al*., 2021; Durand & Robson, 2023; Sellaro *et al*., 2024). Yet, our mechanistic understanding of plant growth responses to neighbour cues comes largely from experiments using steady light conditions that simulate the diffuse background, but do not incorporate LFS. Here we show that LFS perceived by the photo-sensory system can reduce the magnitude of shade avoidance. phyB, cry1, cry2 and UVR8 were sensitive to short, minute-scale stimuli provided by LFS causing persistent hypocotyl growth inhibition (Fig. 1d-g).

### The response to LFS requires synergism among photoreceptors

Laboratory experiments involving different systems have repeatedly demonstrated synergistic co-action among photo-sensory receptors, where some of them enhance the responsivity to others and therefore, all are required for optimum effectiveness (Mohr, 1994; Casal & Mazzella, 1998; Boccalandro *et al*., 2001; Sellaro *et al*., 2009; Fierro *et al*., 2015). Under controlled conditions, this interdependency is conditional because it occurs in plants exposed to limiting light inputs and not under stronger light inputs, where the photoreceptors become mutually redundant (Casal & Mazzella, 1998). However, the scenario (if any) where synergic co-action could operate in natural environments had not been elucidated. Here we show that the synergism among photoreceptors is important for the sensory perception of the sub-optimal (short) light input provided by LFS. In fact, mutating any of the phyB, phyA, cry1 or cry2 photoreceptors was enough to render WL LFS ineffective (Fig. 1d). Similarly, nuclear translocation of UVR8 was stronger under WL+UV-B LFS than UV-B LFS (Fig. 3f), suggesting that other photoreceptors facilitate UVR8 nuclear translocation under short UV-B.

### LFS initiate positive feed-back loops that prime the seedlings to more sensitive responses to light

Repeated LFS render the seedlings sensitive to the weak levels of blue light available under shade, a cue that is not perceived by shade controls not previously exposed to LFS (Fig. 2g). WL LFS increased cry1 nuclear abundance and the sensitivity to blue light (Fig 2e, g). Photo-activated cry1 forms homo-oligomers via its amino terminal domain, which are a necessary step for biological activity (Sang *et al*., 2005). This is a rapid response, predicted to advance significantly during the 2 min of the LFS (Liu *et al*., 2020). cry1 forms well-defined nuclear bodies under blue light (Liu *et al*., 2022), and repeated WL LFS increased the heterogeneity of sub-nuclear localisation consistently with an incipient stage of nuclear body formation (Fig. 2f). Under continuous, high intensity (100 μmol m^-2^ s^-1^) blue light, nuclear cry1 undergoes degradation in the proteasome partially masked by a stable cytoplasmic pool (Liu *et al*., 2022). However, under repeated LFS cry1 accumulated in the nucleus via post-transcriptional mechanisms (Fig. 2e, S4b). Therefore, the abundance of cry1 apparently adjusts to the light input, increasing when it is limiting (e.g., LFS) and decreasing when it is intense.

UV-B exposure during LFS increased the effectiveness of subsequent LFS (Fig. 3b-c). WL+UV-B LFS increased UVR8 nuclear abundance and the response to UV-B (Fig. 3a, d-e). Under prolonged UV-B, UVR8 switches from homodimer to monomer (Rizzini *et al*., 2011), which migrates to the nucleus, apparently via free diffusion, where it accumulates without showing changes in its total protein levels (Kaiserli & Jenkins, 2007; Yin *et al*., 2016; Fang *et al*., 2022). After termination of prolonged UV-B, the monomer slowly back reverts to its homo-dimeric ground state (Heilmann & Jenkins, 2013; Heijde & Ulm, 2013; Podolec *et al*., 2021b) and UVR8 returns from the nucleus to the cytosol (Fang *et al*., 2022). A single LFS was enough to monomerise UVR8, but the monomer weakly accumulated in the nucleus and re-dimerised surprisingly rapidly after the end of the LFS (Fig. 3a-b). Repeated LFS did not increase the proportion of monomer beyond that established by 1 LFS but favoured its nuclear accumulation (without affecting total UVR8 protein levels) and slowed down its re-dimerization. The UVR8 monomer partially returned to the cytosol during the 6 min following the LFS (Fig. 3f), indicating its requirement of UV-B to remain in the nucleus. The interaction of the UVR8 monomer with COP1, which retains UVR8 in the nucleus, is enhanced by UV-B (Liao *et al*., 2020; Fang *et al*., 2022).

### phyB is ready to respond under shade

Photo-activated phyB forms nuclear bodies (Hahm *et al*., 2020) in a process that involves liquid-liquid phase separation (Chen *et al*., 2022). WL LFS did not affect the sub-nuclear distribution of phyB (Fig. 2a), which is consistent with previous results showing no significant changes in the nuclear bodies during the first 4 min after the light to shade or shade to light transitions (Trupkin *et al*., 2014). The duration of the LFS used here (2 min) should be enough to drive substantial photoconversion of the Pr monomer of the inactive Pr-Pfr conformation to generate active Pfr-Pfr dimers (Klose *et al*., 2015; Burgie *et al*., 2021). phyB nuclear bodies were observed under our simulated shade. Thus, phyB would be in a state of facilitation in nuclear condensates, where rapid interaction with partners already present there (Kim *et al*., 2023) would be feasible after transformation to Pfr-Pfr of the Pr-Pfr dimer pool predicted to be present in these nuclear bodies (Klose *et al*., 2015).

### Signalling dynamics downstream the photoreceptors

The hypocotyl growth response to LFS required PIF4 (Fig. 4a) and PIF4 binding targets were overrepresented among the genes with expression reduced by LFS, particularly genes linked to auxin and required for hypocotyl growth under shade (Fig. 5). phyB (Pham *et al*., 2018), cry1, cry2 (De Wit *et al*., 2016; Pedmale *et al*., 2016; Ma *et al*., 2016) and UVR8 (Sharma *et al*., 2019; Tavridou *et al*., 2020b,a) reduce the abundance and/or intrinsic activity of PIF4 and repeated LFS did reduce the abundance of PIF4 (Fig. 4e). PIF4 dynamics contributes to the persistent effects of LFS. In fact, the response to PIF4 nuclear abundance to an interruption of shade was asymmetric, significantly faster in the way down upon exposure to WL+UV-B than during the recovery after returning to shade (Fig. 4e).

The hypocotyl growth response also required COP1 and BES1 (Fig. 4b-c). Although the nuclear levels of COP1 or BES1 did not decrease in response to LFS (Fig. 4f-g), phyB, cry1, cry2 and UVR8 negatively regulate COP1 (Podolec & Ulm, 2018; Ponnu & Hoecker, 2021) and BES1 (Liang *et al*., 2018; Wang *et al*., 2018; Wu *et al*., 2019) by additional molecular mechanisms that could operate under LFS. The hypocotyl growth response to LFS did not require HY5 and nuclear HY5-YFP did not change in response to LFS (Fig. 4d, h). Conversely, HY5 is important for the hypocotyl growth responses to sun patches and sun gaps (Sellaro *et al*., 2011; Moriconi *et al*., 2018), demonstrating fundamental differences in the mechanisms involved in the response to light fluctuations over diverse time scales.

### Functional implications of the response to LFS

For the photosynthetic performance, LFS are a double-edged sword. On the one hand, LFS represent a significant proportion of the light input, and the photosynthetic apparatus can take advantage of them. On the other hand, LFS create a scenario where high light suddenly reaches leaf tissues acclimated to shade, potentially damaging the photosynthetic apparatus (Way & Pearcy, 2012; Kaiser *et al*., 2017; Demmig-Adams *et al*., 2022; Long *et al*., 2022) (Fig. 1a-b). Therefore, the perception of LFS and concomitant reduction in the magnitude of the shade-avoidance response appear to be critical to prevent an excessive exposure and damage of the shade-acclimated photosynthetic apparatus.

LFS perceived by photo-sensory receptors initiated massive changes in the expression of genes involved in the photosynthetic apparatus. Although, within the time frame used here, we observed a negative impact of LFS on the photosynthetic capacity (Fig. 1b), the latter changes of the transcriptome suggest an ongoing process of acclimation to the stronger light input under LFS. Noteworthy, LFS enhanced the expression of genes induced by water deprivation and reduced the impact of later water restriction on growth (Fig. 5). In the field, sunflecks drive the opening of stomata and elevate the temperature of plant tissues increasing vapour pressure deficit (Long *et al*., 2022). These two changes accelerate transpiration and potentially deteriorate water status, suggesting an adaptive value of the observed changes. These findings support the hypothesis that UV-B triggers pre-emptive acclimation to drought (Aphalo & Sadras, 2022). Most plant canopies similarly reduce blue and red wavebands (Morgan *et al*., 1985; Casal, 2012), which activate cry1, cry2 and phyB, but UV-B is more attenuated as it penetrates through the canopy (Durand & Robson, 2023). Thus, perception of UV-B by UVR8 would inform the nearness to the edge of the canopy, enhancing the negative correlation between shade avoidance and light exposure.

## Conclusions

LFS reduce shade avoidance thanks to the amplification of the light signal by the sensory system. First, multiple photosensory receptors, which tend to be redundant under strong light inputs operate synergistically under the weak input of LFS. Second, both cry1 and UVR8 increase their nuclear abundance in response to repeated LFS and gain sensibility to light. Positive feedback motifs are crucial to allow for the buffering of propagated noise caused by rapid input fluctuations while maintaining sensitivity to long-term changes in the input signal (Hornung & Barkai, 2008). Third, the perception of LFS reduces the nuclear abundance of PIF4 to curb shade avoidance, and PIF4 is slow to revert this decay after the impact of light. The implications of sensory perception of LFS go beyond preventing the exposure of shade acclimated tissues to excessive light to trigger transcriptional responses of photosynthetic and water restriction genes, which adjust the seedlings to the environmental opportunity and challenges generated by the presence of LFS.

## Acknowledgements

We thank Chentao Lin, Qin Wang and Xu Wang (Fujian Agriculture and Forestry University, China) for kindly providing seeds of the transgenic lines expressing cry1 or cry2 fused to GFP, and Belen Borniego and Cristian Escudero (University of Buenos Aires and CONICET) for help with genotyping. This research was supported by a grant from the Argentinean-Swiss Joint Research Programme of CONICET-MINCyT-SNSF (grant no. IZSAZ3_173361 to R.U. and J.J.C.).

## Author contributions

R.U. and J.J.C designed the study; A.B., N.T., E.P. and C.C. performed the experiments; A.B., A.R. and J.J.C. analysed the data; and J.J.C. wrote the paper with input from A.B. and R.U.

## Competing interests

The authors declare no competing interests.

## Supplementary information

**Fig. S1.**
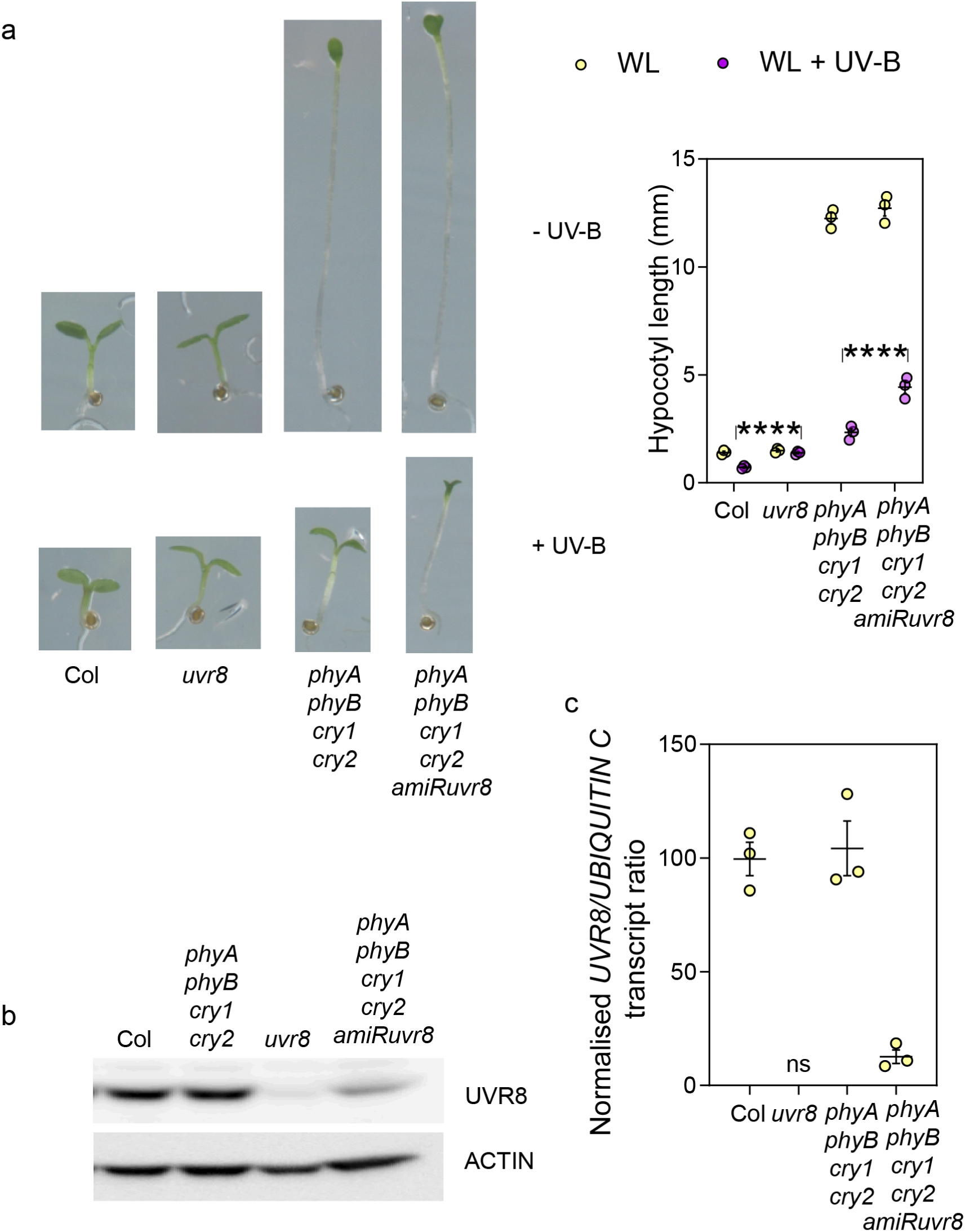
Basic characterisation of the *phyA phyB cry1 cry2 amiRuvr8* line. (a) Hypocotyl length of seedlings of the *phyA phyB cry1 cry2* quadruple mutant and the *phyA phyB cry1 cry2 amiRuvr8* line under continuous WL with or without supplementary UV-B and photographs of representative seedlings. (b) UVR8 protein levels, ACTIN was used as loading control. **c** Transcript levels of *UVR8* relative to *UBIQUITIN C*, normalised to the values observed in the wild type. The seedlings were grown for five days under continuous white light (a-c, 10 µmol m^-2^ s^-1^) or white light plus UV-B (a, 1.5 µmol m^-2^ s^-1^ provided by Philips TL20W/01RS narrowband UV-B tubes) before the measurements. Data are means and ±SE of three biological replicates (ten seedlings per replicate). Asterisks indicate significant differences (****, P <0.0001) between the indicated means. The primers were *UBQsen*: TTCCTTGATGATGCTTGCTC, *UBQant:* TTGACAGCTCTTGGGTGAAG, *UVR8sen:* GGTAAATGAAATGGTCAAGAAACAAA, *UVR8ant:* AATACGGACCTCACCGGTAAAG.

## Notes

### Competing Interest Statement

The authors have declared no competing interest.

### Summary of Updates

Ddditional controls were added, and the writing format has been improved for better clarity and compliance with standard guidelines

