## Supplementary material for "Sensory perception of fluctuating light in Arabidopsis": Suplemental material

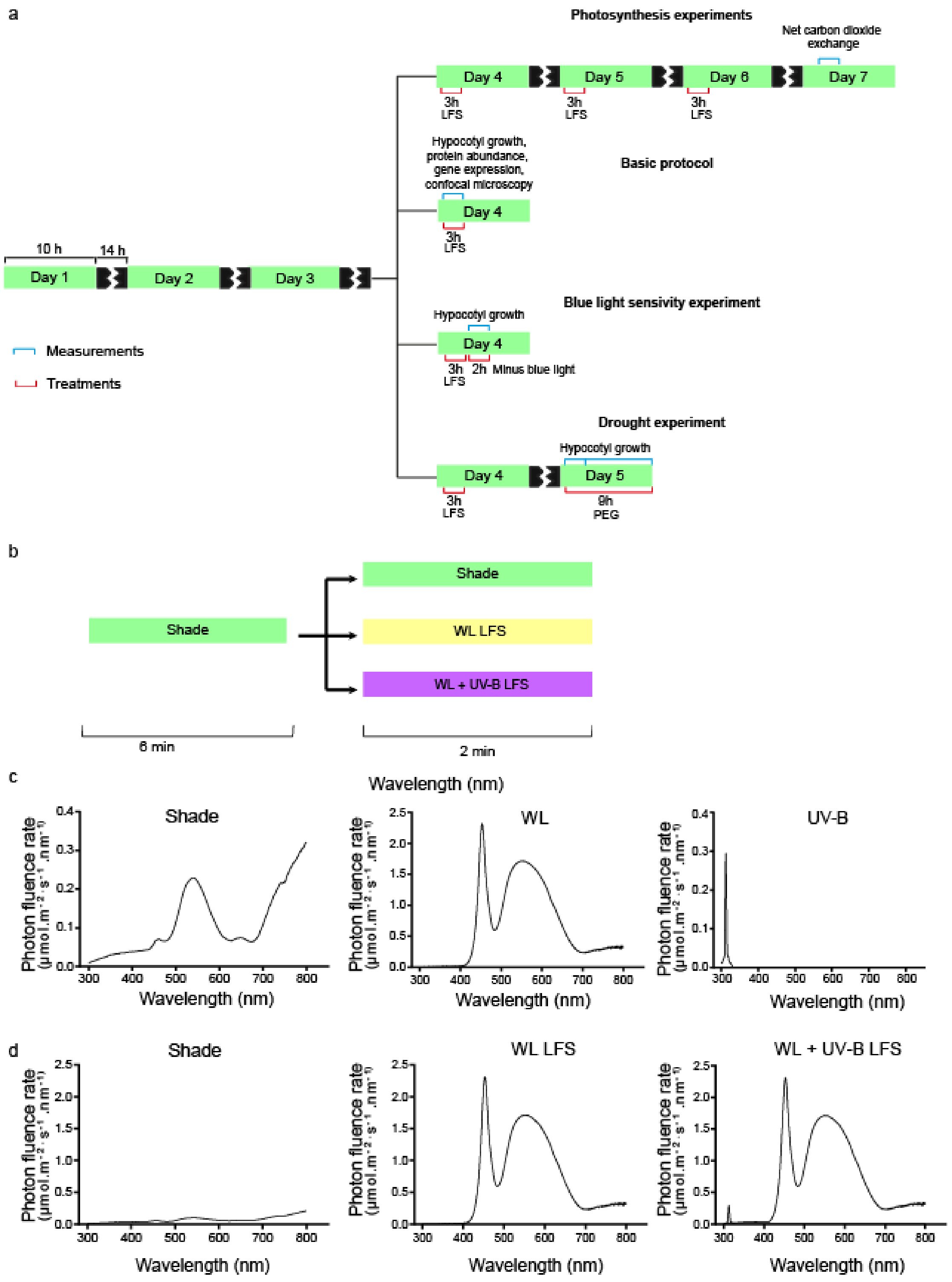


**Fig. S2.** Experimental protocols. (a) Protocol used in growth, confocal microscopy, protein blots, gene expression and photosynthesis experiments. (b) During the 3 h of treatment, the seedlings were exposed to pulses of white light (WL) or WL + UV-B (WL + UV-B) for 2 min followed by 6 min shade (completing a LFS cycle every 8 min) and the rest of the photoperiod remained under shade. (c-d) Spectral photon distribution provided by each light source (c, note different scales) and experimental condition (d, note the same scale), measured with an Ocean Optics spectroradiometer.

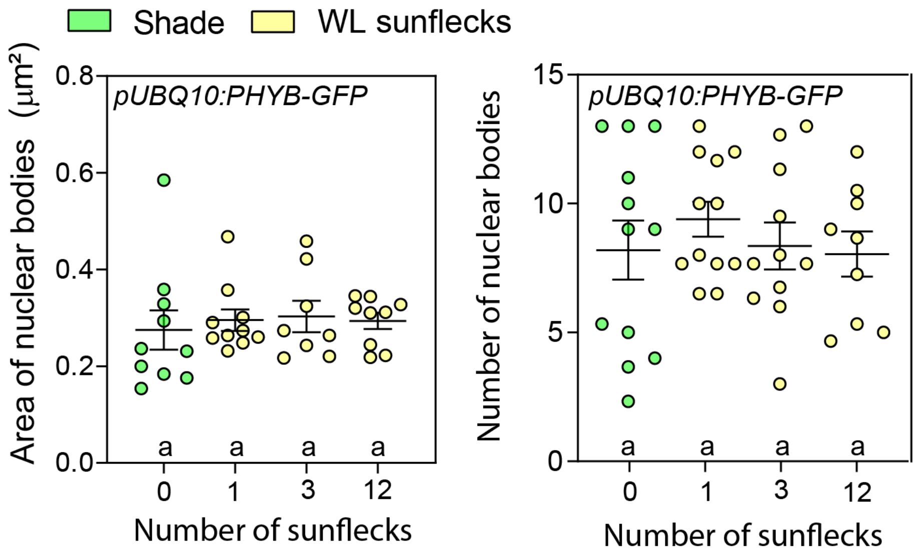


**Fig. S3.** LFS do not affect the size or number of phyB nuclear bodies. Data are means ±SE and individual values of 8-12 biological replicates. Similar letters indicate the absence of significant differences in Tukey multiple comparison test.

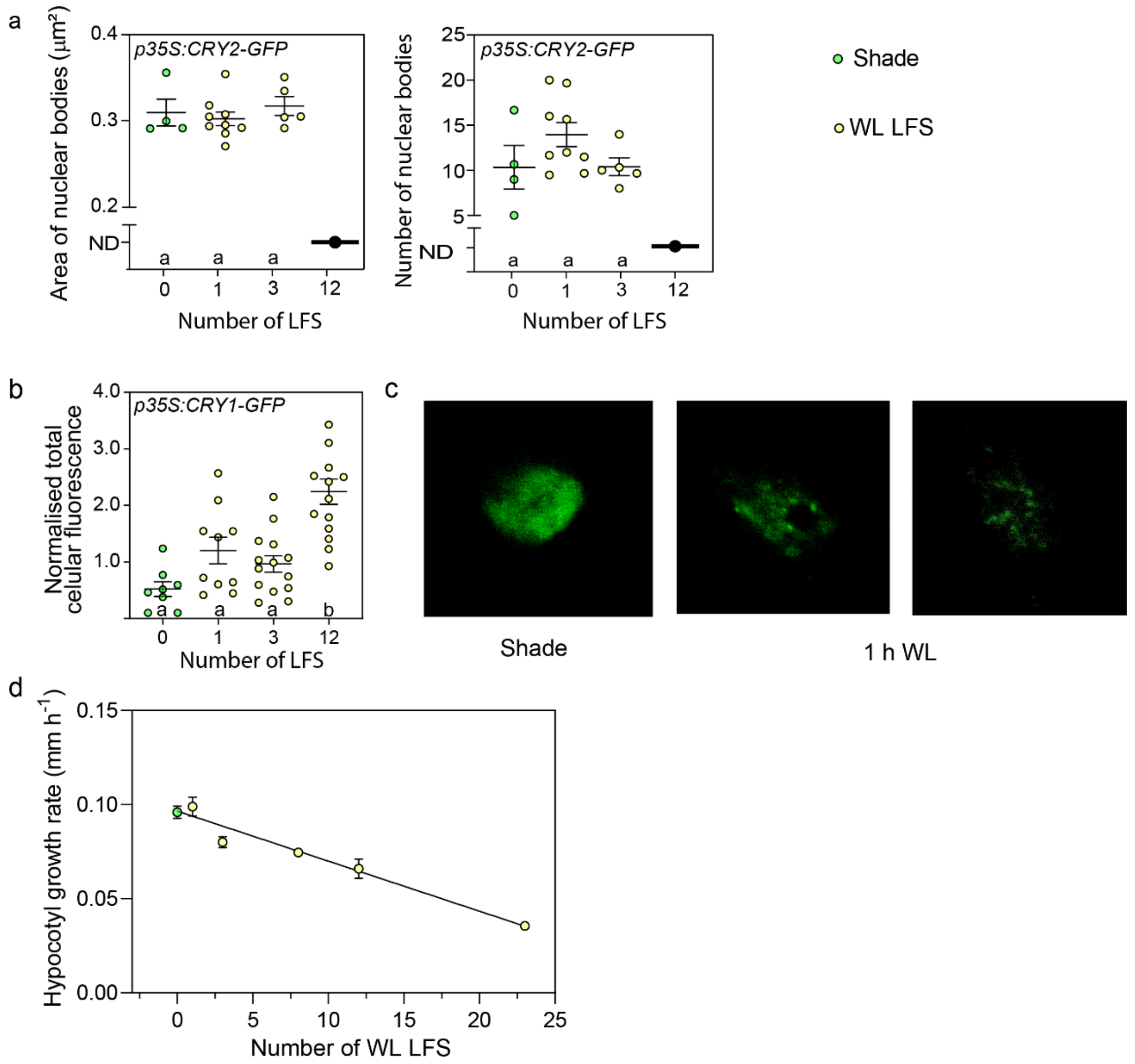


**Fig. S4.** Dynamics of cry2, cry1 and response to WL LFS. (a) LFS reduce the size and number of cry2-GFP nuclear bodies. ND, not detectable. (b) Total cellular fluorescence driven by cry1-GFP increases after 12 LFS. (c) cry1-GFP forms nuclear bodies in seedlings grown under shade and exposed to 1 h continuous white light. (d) Hypocotyl growth decreased linearly with the number of WL LFS (varied by changing their frequency). This indicates a constant contribution of each LFS. Data are means ±SE (a-b, d) and individual values (a-b) of 4-9 biological replicates. Significant differences in Tukey's multiple tests indicated by different letters ($P < 0.05$). The linear regression (d) is significant at $P < 0.001$.

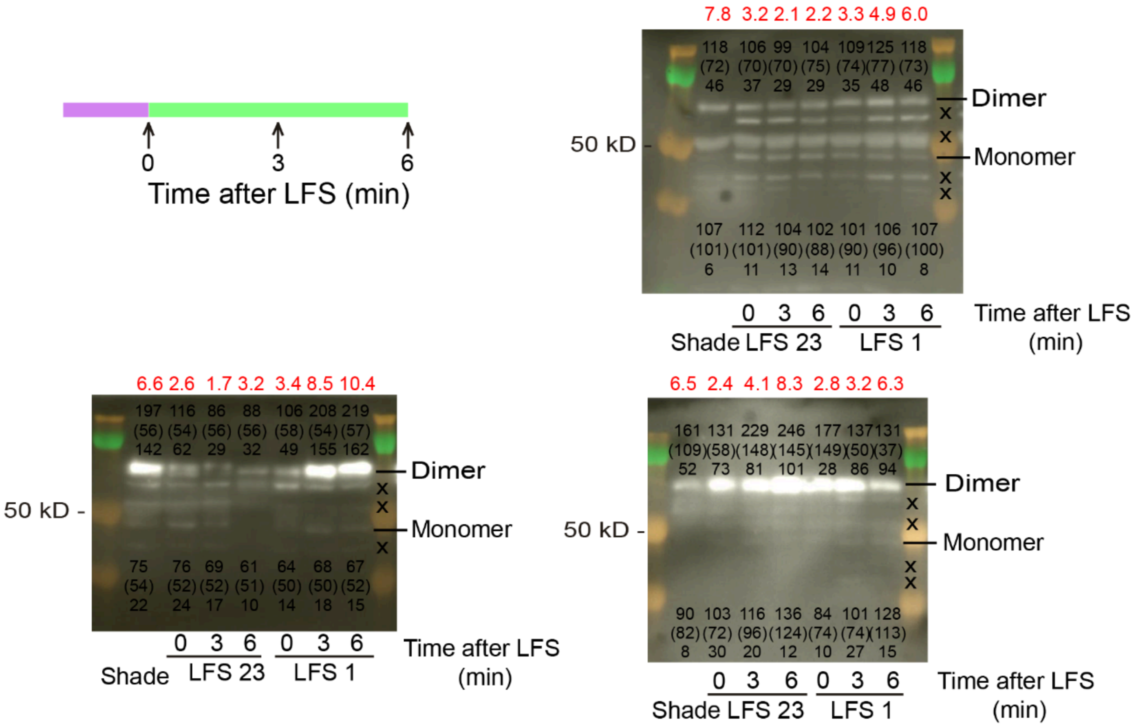


**Fig. S5.** Quantification of dimer / monomer ratios in UVR8 protein blots. The blots correspond to different biological replicates (data summarised in Fig. 3). "x" unspecific bands. The bands corresponding to the UVR8 dimer and monomer were revealed with anti-UVR8$^{(426\text{-}440)}$ (Favory *et al.*, 2009) (shown here) and confirmed with a different antibody (anti-UVR8$^{(410\text{-}424)}$) (Heijde & Ulm, 2013). For dimer and monomer bands we show the intensity, the intensity of its background (in brackets), and the difference between band and background intensity. The ratio between dimer and monomer is indicated at the top of the blot (in red).

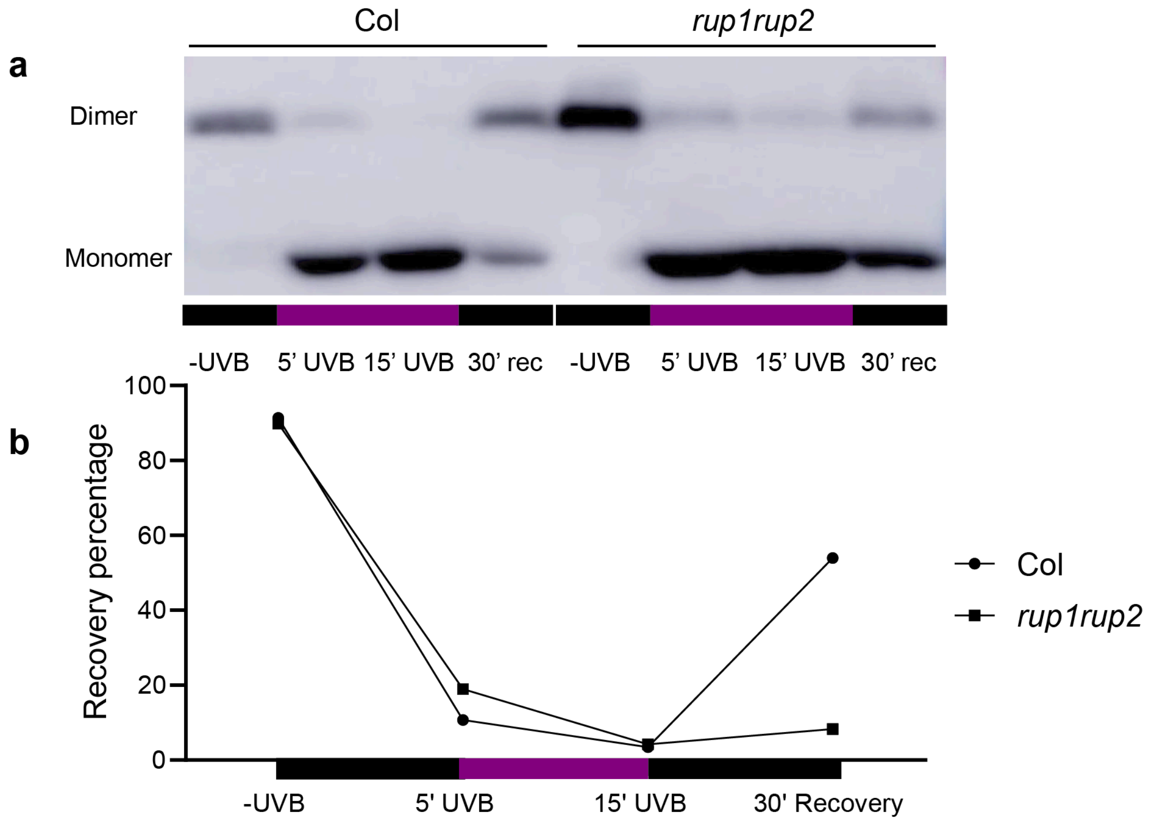


**Fig. S6.** Slow re-dimerization of UVR8 in darkness after the exposure to prolonged UV-B. (a) UVR8 dimer-monomer dynamic in Col and *rup1rup2* mutant. (b) Recovery percentage of UVR8 dimer after 30 minutes of darkness after 0, 5 or 15 minutes of UVB exposure.

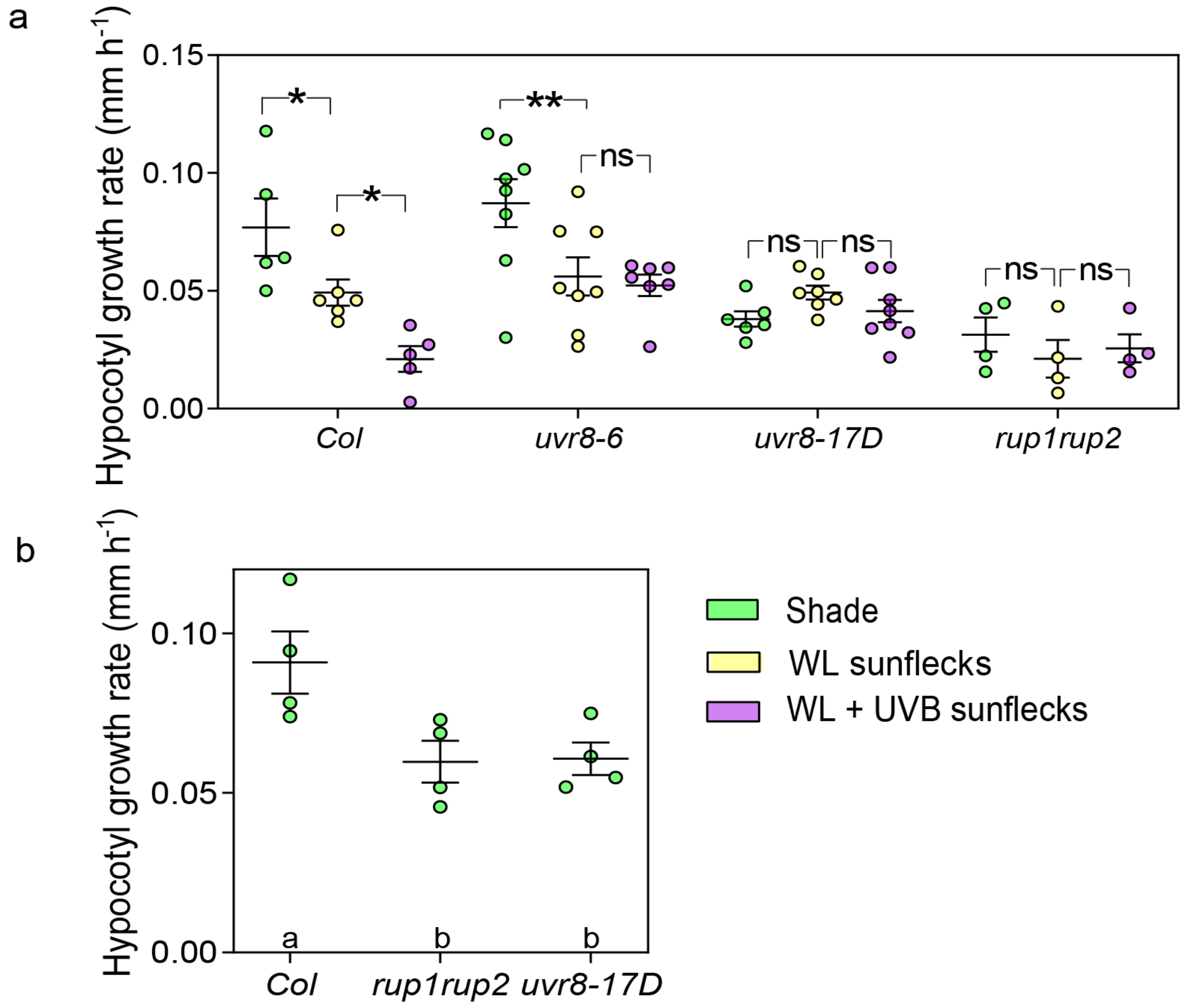


**Fig. S7.** The perception of LFS requires re-dimerization of UVR8. (a) The *uvr8-17D* and *rup1 rup2* mutants impaired in UVR8 dimerization fail to respond to UV-B. (b) The *uvr8-17D* and *rup1 rup2* mutants are shorter than the wild type even under shade, indicating that residual UV-B under this light condition (Fig. S1c) is enough to establish saturating levels of UVR8 activity in these mutants. The experiments in (b) were conducted in the absence of sunflecks to avoid any unnoticed WL or UV-B coming from LFS treatments. Data are means ±SE and individual values of 4-8 biological replicates. Significant differences in Tukey's multiple tests indicated by different letters ($P < 0.05$) or by asterisks (*, $P < 0.05$; **, $P < 0.01$, ns, not significant).

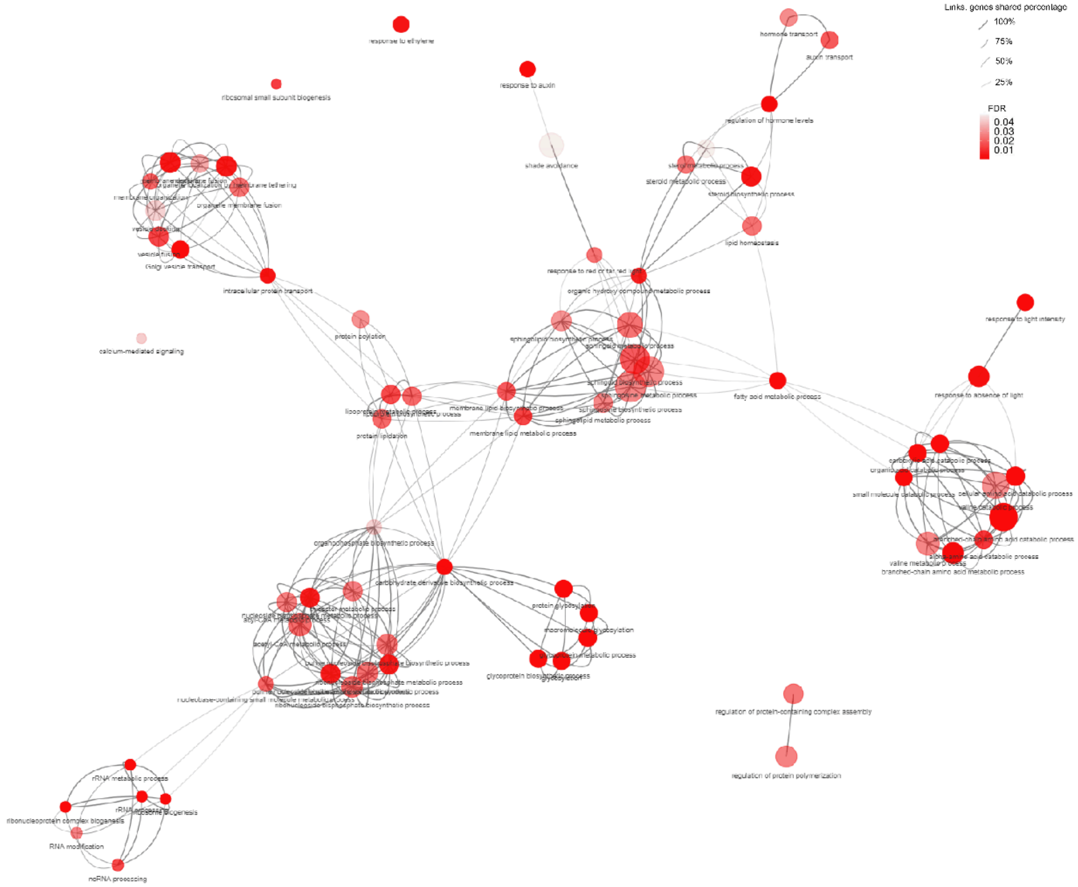


**Fig. S8.** GO terms overrepresented among the genes with expression reduced by LFS in a cry1 cry2 and UVR8-dependent manner (Cluster 2). The size of each node is determined by fold enrichment, node colour is indicative of the false discovery rate, and the width of connecting edges represents the number of shared genes.

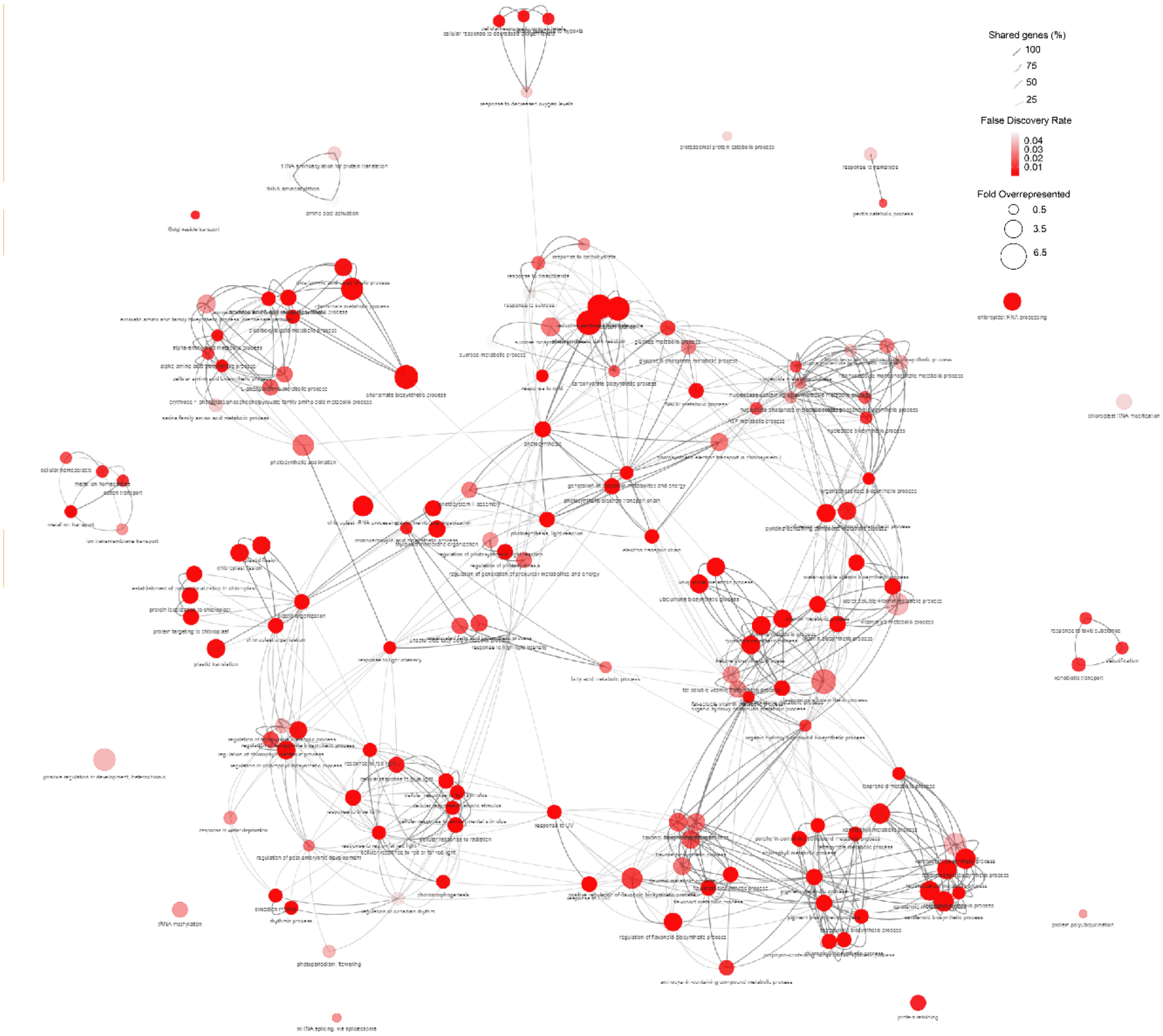


**Fig. S9.** GO terms overrepresented among the genes with expression enhanced by LFS in a cry1 cry2 and UVR8-dependent manner (Cluster 1). The size of each node is determined by fold enrichment, node colour is indicative of the false discovery rate, and the width of connecting edges represents the percentage of shared genes.

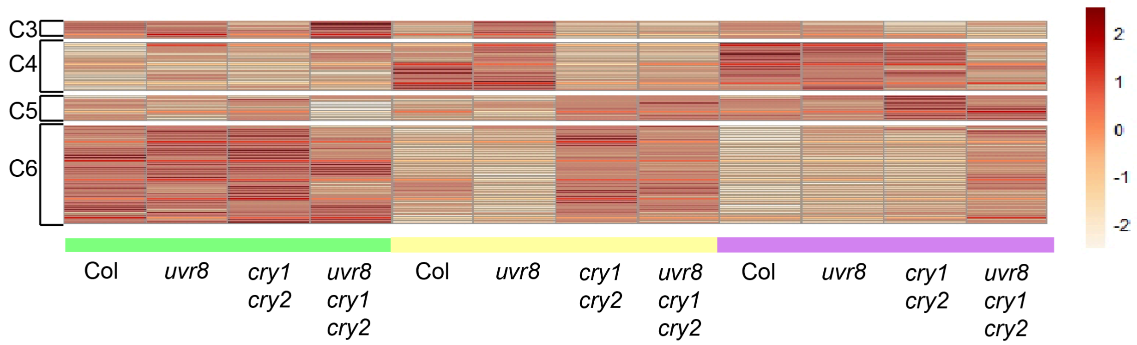


**Fig. S10.** Genes that responded to LFS in a cry1 cry2 dependent manner but independently of UVR8. Heatmap of the gene clusters 3-6 (C3-C6). No GO terms were significantly overrepresented in these clusters.

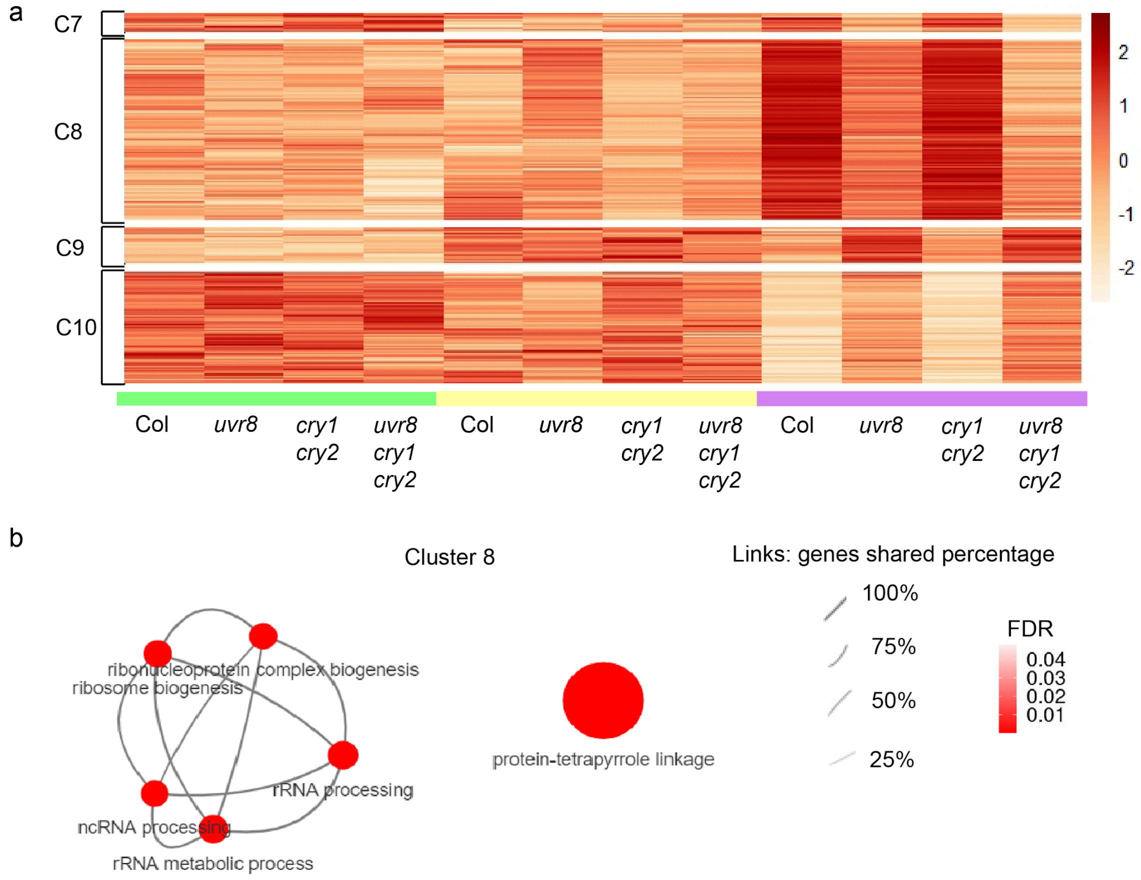


**Fig. S11.** Genes that responded to LFS in a UVR8 dependent manner but independently of cry1 cry2. (a) Heatmap of the gene clusters 7-10 (C7-10). (b) GO terms overrepresented in cluster 8. Only "regulation of cellular ketone metabolic processes was overrepresented in cluster 9 (corrected P <0.04) and no GO terms were significantly overrepresented in clusters 7 and 10.

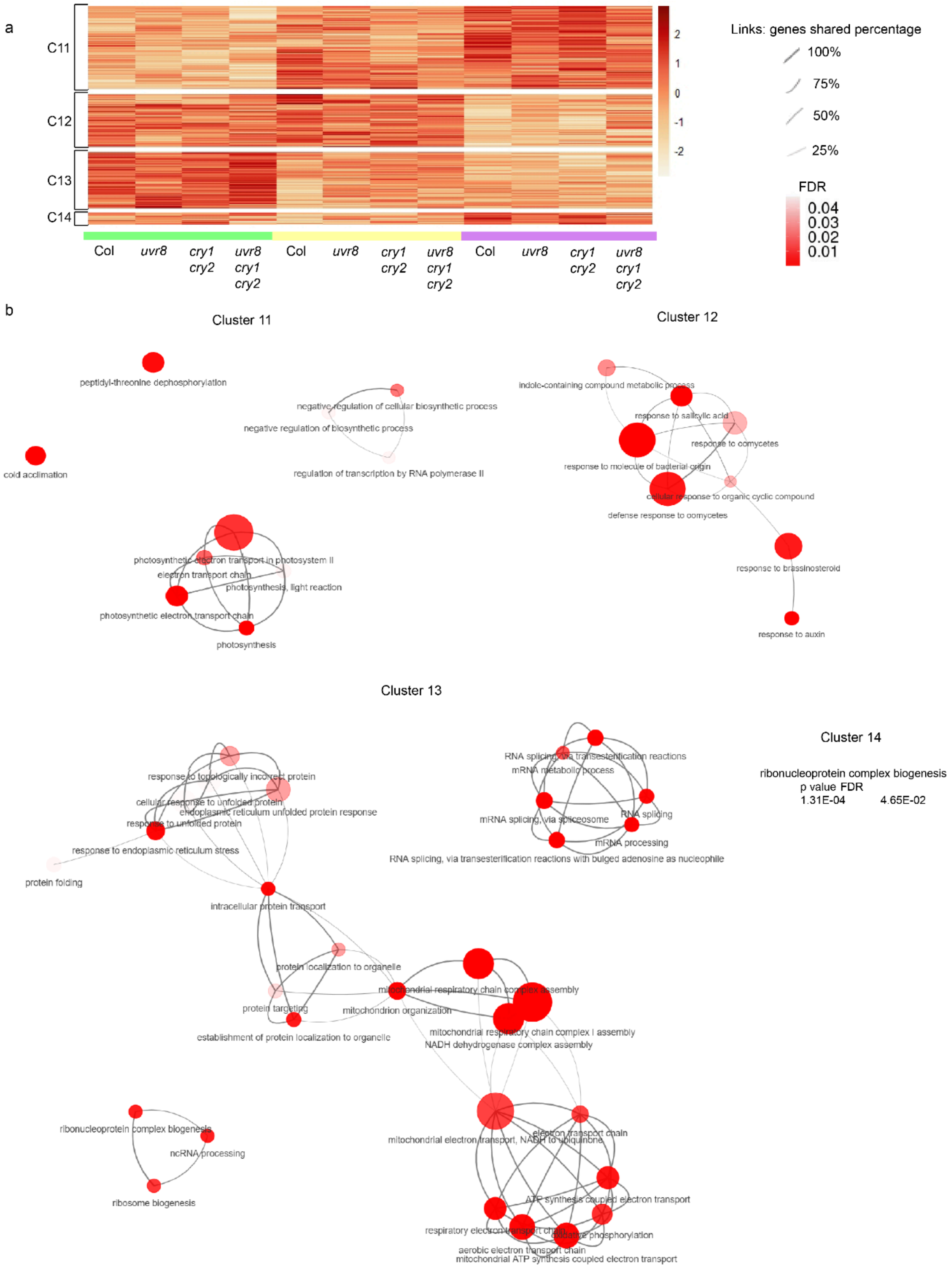
a
C11
C12
C13
C14
2
1
0
-1
-2
Links: genes shared percentage
100%
75%
50%
25%
FDR
0.04
0.03
0.02
0.01
Col
uvr8
cry1 cry2
uvr8 cry1 cry2
Col
uvr8
cry1 cry2
uvr8 cry1 cry2
Col
uvr8
cry1 cry2
uvr8 cry1 cry2
b
Cluster 11
peptidyl-threonine dephosphorylation
cold acclimation
negative regulation of cellular biosynthetic process
negative regulation of biosynthetic process
regulation of transcription by RNA polymerase II
photosynthetic electron transport in photosystem II
electron transport chain
photosynthesis, light reaction
photosynthetic electron transport chain
photosynthesis
Cluster 12
indole-containing compound metabolic process
response to salicylic acid
response to oomycetes
response to molecule of bacterial origin
cellular response to organic cyclic compound
defense response to oomycetes
response to brassinosteroid
response to auxin
Cluster 13
response to topologically incorrect protein
cellular response to unfolded protein
endoplasmic reticulum unfolded protein response
response to unfolded protein
response to endoplasmic reticulum stress
protein folding
intracellular protein transport
protein localization to organelle
mitochondrial respiratory chain complex assembly
protein targeting
mitochondrion organization
establishment of protein localization to organelle
mitochondrial respiratory chain complex I assembly
NADH dehydrogenase complex assembly
RNA splicing, via transesterification reactions
mRNA metabolic process
RNA splicing
mRNA splicing, via spliceosome
mRNA processing
RNA splicing, via transesterification reactions with bulged adenosine as nucleophile
ribonucleoprotein complex biogenesis
ncRNA processing
ribosome biogenesis
mitochondrial electron transport, NADH to ubiquinone
electron transport chain
ATP synthesis coupled electron transport
oxidative phosphorylation
aerobic electron transport chain
mitochondrial ATP synthesis coupled electron transport
Cluster 14
ribonucleoprotein complex biogenesis
p value FDR
1.31E-04 4.65E-02

**Fig. S12.** Genes that responded to LFS independently of cry1 cry2 and UVR8. (a) Heatmap of the gene clusters 11-14 (C11-C14). (b) GO terms overrepresented in clusters 11-13. Only "ribonucleoprotein complex biogenesis" (corrected P <0.05) was overrepresented in cluster 14.

Table S1. Mutant and transgenic lines used in this study.

| Mutant lines | Reference |
|---|---|
| *bes1-1* | (He *et al.*, 2005) |
| *cop1-4* | (McNellis *et al.*, 1994) |
| *cry1-304* | (Mockler *et al.*, 1999) |
| *cry1-304 cry2-1* | (Mockler *et al.*, 1999) |
| *cry2-1* | (Mockler *et al.*, 1999) |
| *hy5-221* | (Shin *et al.*, 2007) |
| *iaa19-1* (CS25217) | This report |
| *iaa29* (SALK_091933) | (Sun *et al.*, 2013) |
| *phyA-211* | (Reed *et al.*, 1994) |
| *phyA* (Salk line N520360) *phyB* (Salk line N569700) | (Zhang *et al.*, 2017) |
| *phyA-412 phyB-9 cry1-304 cry2-1* | (Strasser *et al.*, 2009) |
| *phyA-412 phyB-9 cry1-304 cry2-1 amiRuvr8* | This report |
| *phyB-9* | (Reed *et al.*, 1993) |
| *pif4-101* | (Lorrain *et al.*, 2008) |
| *pin3-3* | (Keuskamp *et al.*, 2010) |
| *pin7-1* | (Keuskamp *et al.*, 2010) |
| *rup1-1 rup2-1* | (Gruber *et al.*, 2010) |
| *uvr8-6* | (Favory *et al.*, 2009) |
| *uvr8-17D* | (Podolec *et al.*, 2021) |
| *cry1-304 cry2-1 uvr8-6* | This report |

| Transgenic lines | Reference |
| --- | --- |
| *uvr8-6 pUVR8:YFP-UVR8* | (Bernula *et al.*, 2017) |
| *uvr8-6 p35S:YFP-UVR8* | (Heijde & Ulm, 2013) |
| *phyB-9 pUBQ10:PHYB-YFP* | (Zhang *et al.*, 2013) |
| *cry1-304 cry2-1 35S:CRY2-GFP* | (Yu *et al.*, 2009) |
| *cry1-304 35S:GFP-CRY1* | (He *et al.*, 2019) |
| *pBES1:BES1-GFP* | (Yin *et al.*, 2002) |
| *cop1-4 pCOP1:myc-mCHERRY-COP1* | (Costigliolo Rojas *et al.*, 2022) |
| *pif4-101 pPIF4:PIF4-GFP* | (Pucciariello *et al.*, 2018) |
| *hy5-1 pHY5:HY5-YFP* | (Oravecz *et al.*, 2006) |

Table S2. Explanatory variables tested in the step-wise multiple regression analysis used for Fig. 1d. The response variable was hypocotyl growth rate$^{-1}$. (a-b), Variables that account for effects of WL or UV-B not mediated by the photoreceptors included in the analysis. The values assumed by WL were shade= 0, WL LFS= 1, WL+UV-B LFS=1. The values assumed by UV-B were shade= 0, WL LFS= 0, WL+UV-B LFS=1. **c**, Variable that accounts for the reported effects of phyA under shade(Yanovsky *et al.*, 1995). The values assumed by phyA were wild type allele= 1, mutant allele= 0. (d-r)**,** Variables that represent the interactions among WL and phyA, phyB, cry1 and/or cry2. The values assumed by these interactions were the products of the values corresponding to WL for that condition by the number corresponding to the wild type or mutant allele of the indicated photoreceptor(s). (s), Variable that represents the interaction between UV-B and UVR8. The values followed the same criteria used for (d-r).

| | **variable** | **excluded/retained** | **slope/constant ± SE** | **p-value** |
|---|---|---|---|---|
| (a) | WL | excluded | - | - |
| (b) | UV-B | excluded | - | - |
| (c) | phyA | retained | 14.34 ± 3.29 | <0.0001 |
| (d) | WL x cry1 | excluded | - | - |
| (e) | WL x cry1 x cry2 | excluded | - | - |
| (f) | WL x cry1 x cry2 x phyA | excluded | - | - |
| (g) | WL x cry1 x phyA | excluded | - | - |
| (h) | WL x cry2 | excluded | - | - |
| (i) | WL x cry2 x phyA | excluded | - | - |
| (j) | WL x phyA | excluded | - | - |
| (k) | WL x phyB | excluded | - | - |
| (l) | WL x phyB x cry1 | excluded | - | - |
| (m) | WL x phyB x cry1 x cry2 | retained | 7.74 ± 2,46 | 0.0018 |
| (n) | WL x phyB x cry2 | excluded | - | - |
| (o) | WL x phyB x phyA | excluded | - | - |
| (p) | WL x phyB x phyA x cry1 | excluded | - | - |
| (q) | WL x phyB x phyA x cry1 x cry2 | excluded | - | - |
| (r) | WL x phyB x phyA x cry2 | excluded | - | - |
| (s) | UV-B x uvr8 | retained | 16.29 ± 2.60 | <0.0001 |
| (t) | constant | - | 3.49 ± 3.35 | 0.2988 |
